# Nebulized delivery of a broadly neutralizing SARS-CoV-2 RBD-specific nanobody prevents clinical, virological and pathological disease in a Syrian hamster model of COVID-19

**DOI:** 10.1101/2021.11.10.468147

**Authors:** Thomas J. Esparza, Yaozong Chen, Negin P. Martin, Helle Bielefeldt-Ohmann, Richard A. Bowen, William D. Tolbert, Marzena Pazgier, David L. Brody

## Abstract

There remains an unmet need for globally deployable, low-cost therapeutics for the ongoing severe acute respiratory syndrome coronavirus 2 (SARS-CoV-2) pandemic. Previously, we reported on the isolation and *in vitro* characterization of a potent single-domain nanobody, NIH-CoVnb-112, specific for the receptor binding domain (RBD) of SARS-CoV-2. Here, we report on the molecular basis for the observed broad *in vitro* neutralization capability of NIH-CoVnb-112 against variant SARS-CoV-2 pseudoviruses, including the currently dominant Delta variant. The structure of NIH-CoVnb-112 bound to SARS-CoV-2 RBD reveals a large contact surface area overlapping the angiotensin converting enzyme 2 (ACE2) binding site, which is largely unencumbered by the common RBD mutations. In an *in vivo* pilot study, we demonstrate effective reductions in weight loss, viral burden, and lung pathology in a Syrian hamster model of COVID-19 following nebulized delivery of NIH-CoVnb-112. These findings support the further development of NIH-CoVnb-112 as a potential adjunct preventative therapeutic for the treatment of SARS-CoV-2 infection.

## INTRODUCTION

The global community continues to face substantial morbidity and mortality due to ongoing coronavirus disease 2019 (COVID-19) caused by the SARS-CoV-2.^1,2^ To date, more than 200 million people have sustained infection and over 4 million have succumbed to the disease worldwide.^3^ The unprecedented effort to develop and bring to the world effective vaccines has proven remarkably successful, yet there remains an unmet need for additional therapeutic options. Notably, antibody treatment from convalescent patient-derived plasma or with recombinant monoclonal antibodies delivered intravenously have demonstrated efficacy in acute infection.^4,5^ While these treatments have potent neutralizing capacity, they come with the limitation of moderate to high cost and intravenous infusion requirement, which may limit their utility in global distribution. This presents an opportunity to explore lower-cost adjunct therapeutics that can be delivered easily. For example, delivery of inhaled treatments to the respiratory system may result in lower dose requirements and direct engagement of the virus at the site of early infection.

The *Camelidae* family, which includes dromedaries, camels, IIamas, and alpacas, possess a subclass of immunoglobulins which consist of paired heavy chains with antigen-binding variable domains.^6^ These single-domain monovalent antigen binding domains, also known as nanobodies, are ∼12 to 15 kDa in size yet still exhibit potent affinity to target antigens.^7^ Moreover, nanobodies can readily be expressed in different expression systems, including bacteria, yeast and many mammalian cell lines.^7,8^ Due to their exceptional stability in a wide range of pH and temperature and their ability to be lyophilized and aerosolized, nanobody therapies have been developed for nebulization delivery in treating respiratory diseases.^10,11^ Nanobodies with potent neutralizing activity against SARS-CoV-2 have been identified by several approaches, such as the immunization of llamas, alpacas and camelid mice with SARS-CoV-2 spike or RBD, screening phage or yeast display libraries of naïve llama nanobody or humanized synthetic nanobody libraries.^9–16^ We previously reported on the isolation and biophysical characterization of a SARS-CoV-2 RBD binding nanobody, NIH-CoVnb-112, which possesses robust biophysical properties and binds to RBD with a low nanomolar affinity.^9^ Thus, nanobodies are an attractive platform for the future development of novel therapeutic, diagnostic, and detection applications.

Here, in order to describe the molecular basis of the neutralization potency and breadth of CoVnb-112, we determined the high-resolution structure of its complex with the SARS-CoV-2 RBD antigen. Our studies reveal that the epitope of NIH-CoVnb-112 largely overlaps the receptor-binding motif that engages the angiotensin converting enzyme 2 (ACE2), which serves as the primary receptor for viral attachment and entry^17,18^ confirming the RBD-specificity of CoVnb-112. Furthermore, structural analyses indicate that recurrent RBD escape mutations are well accommodated within the nanobody-RBD interface with minor disruption of nanobody-RBD contact residues. This modeling data corroborates NIH-CoVnb-112 binding and pseudovirus neutralization assays demonstrate a broad range of SARS-CoV-2 variant data to RBD mutants and *in vitro* neutralization data against pseudotyped SARS-CoV-2 virus from four variants of concern, including the currently dominant Delta (B.1.617.2). Finally, we demonstrate a robust reduction of viral burden and prevention of lung pathology in a hamster model of COVID-19 following nebulized delivery of NIH-CoVnb-112. These results support the case for further development of nanobody-based therapeutics for COVID-19.

## RESULTS

A key advantage of nanobodies is the ability to easily transition to alternative expression systems for scaled material production. Previously, we cloned the NIH-CoVnb-112 coding sequence into the pICZα *Pichia pastoris* expression vector.^9^ Using the same clone, fermentation was performed under defined parameters in a 4-liter DCI Tryton bioreactor. Following 72 hours of monitored feed methanol growth, the supernatant was harvested, concentrated by tangential flow filtration, and purified by immobilized metal affinity chromatography. The final yield for the 4-liter expression was 1.1 grams as measured by absorbance, with a purity of >95%. This preparation was used for all experiments reported here.

### Structural basis of NIH-CoVnb-112 binding to SARS-CoV-2 RBD

To elucidate the broad-spectrum reactivity of NIH-CoVnb-112 against the circulating SARS-CoV-2 variants and to map its epitope, we determined its crystal structure bound to the prototype SARS-CoV-2 RBD at 2.82 Å resolution (**Fig. 1, Fig. 2** and **Supplementary Table 1**). NIH-CoVnb-112 binds to the RBD through a canonical site of vulnerability that significantly overlaps the ACE2 binding site (**Fig. 1a-b**). Over 70% of the contact surface is mediated through residues within the elongated tyrosine-rich Complementarity-Determining Region 3 (CDR3) that forms an extensive shape-complementarity interaction with the receptor binding motif (RBM, residues 403-505) of the RBD (**Fig. 1c-e**). The CDR3 loop traverses the amphiphilic groove flanked by the protruding RBD tip (residues 470-492) and the RBD-core, which is stabilized by an atypical inter-CDR disulfide (C^50^_CDR2_ –C^100H^_CDR3_, as defined using Kabat numbering^19^). This disulfide is conserved in two published nanobody homologs, WNb-2^13^ and VHH-E, isolated from alpaca^10^, both of which also bind the RBD in a similar manner, highlighting the functional importance of this CDR-stabilizing disulfide. The 21 amino acid length CDR3 contains six conserved tyrosines; of these, three pairs of π- π interactions appear to be crucial for the CDR3 loop conformation, including the face-to-face stacking of F^490^_RBD_ and Y^100E^_nb112_ and the two face-to-edge contacts between Y^449^_RBD_-Y^99^_nb112_ and Y^505^_RBD_-Y^102^_nb112_, as documented by their high buried surface area (BSA) values (**Fig. 1d**).

**Figure 1.**
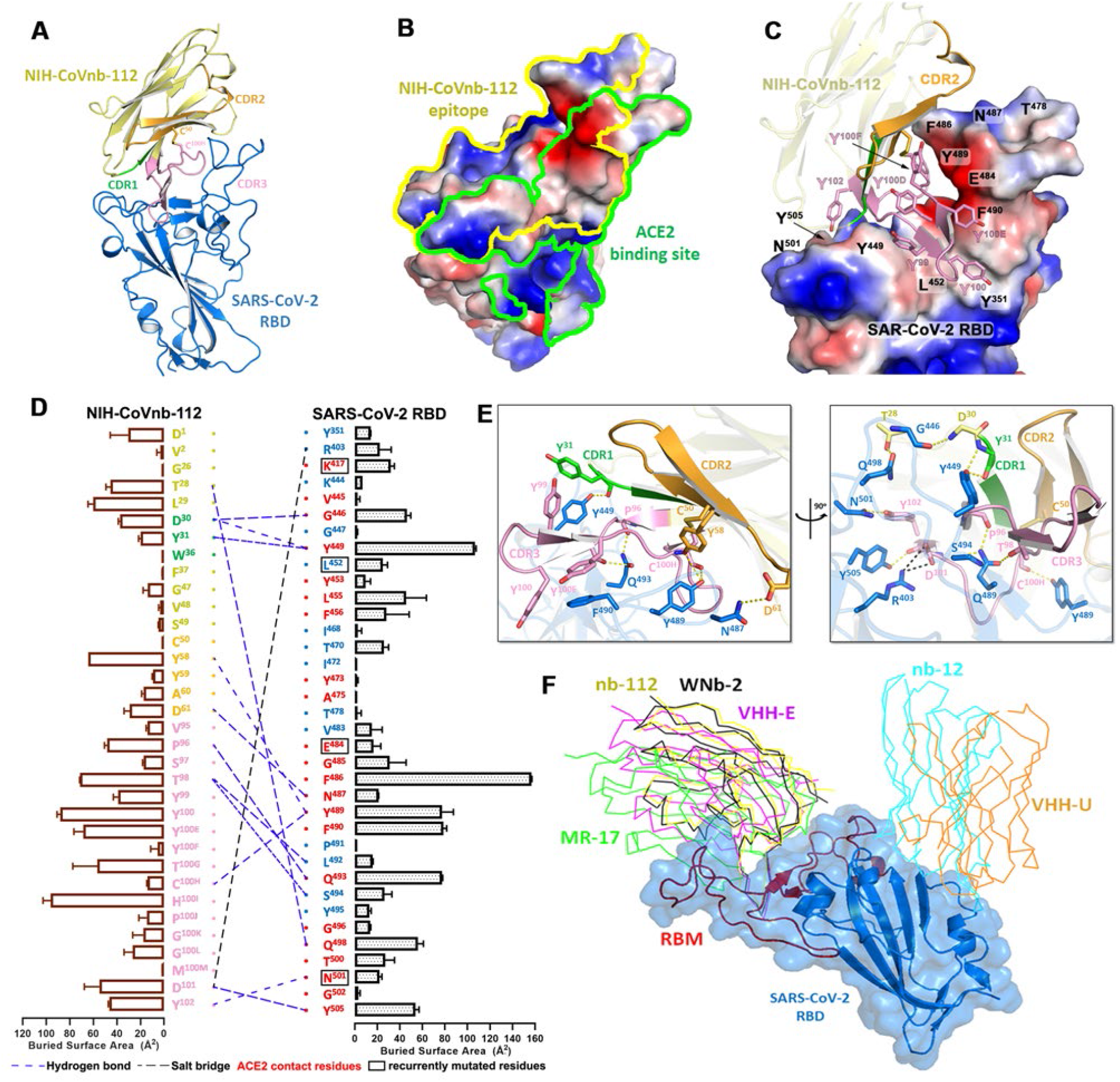
Molecular basis of SARS-CoV-2 spike recognition by NIH-CoVnb-112. (**a**) Crystal structure of NIH-CoVnb-112:SARS-CoV-2 RBD complex. The framework of NIH-CoVnb-112 is shown as yellow ribbons with the CDR1, CDR2, CDR3 colored green, orange and pink, respectively. The SARS-CoV-2 RBD is colored blue. (**b**) Comparison of the RBD-contact footprint of NIH-CoVnb-112 and ACE2. The contact surface area is indicated by yellow and green lines for NIH-CoVnb-112 and ACE2, respectively, over the molecular surface of RBD. The electrostatic potential is displayed over the RBD surface and colored for negative (red), positive (blue), and neutral (white) electrostatic potential, respectively. (**c**) Enlarged view into the NIH-CoVnb-112:RBD interface. NIH-CoVnb-112 is shown as a ribbon with the same color scheme as (**a**) and the SARS-CoV-2 RBD is displayed as an electrostatic potential surface. (**d**) The interaction network at the NIH-CoVnb-112:RBD interface. The individual nanobody:RBD contacts are shown as lines with the diagram of buried surface area for individual interface residue shown on the sides. Nanobody residues in the framework and CDRs are colored as in (**a**) with interactions defined by a 5-Å distance criterion cutoff shown as lines. The Kabat^19^ scheme was used to number nanobody amino acid residues, with unique insertion residues indicated by letter (e.g., 52a, 52b, 52c). Salt bridges and H-bonds (bond length less than 3.5 Å) are shown as black and blue dashed lines, respectively. The ACE2 contact residues within the nanobody epitope are highlighted in red or otherwise in blue. The recurrently mutated RBD residues in SARS-CoV-2 variants are marked with black boxes. (**e**) Orthogonal views of the interaction network between NIH-CoVnb-112 and RBD. CDR residues are colored as in (**a**) and the RBD residues are colored in blue. Hydrogen bonds and salt bridges are denoted as yellow and black broken lines, respectively. (**f**) Comparison of the mode of RBD recognition by NIH-CoVnb-112 to other RBD-specific nanobodies. The NIH-CoVnb-112-RBD complex was superimposed (based on the RBD) to five published RBD complex structures of RBD-specific nanobodies, including WNb-2 (PDB: 7LDJ), VHH-E (7KN5), MR17 (7CAN), Nb12 (7MY3), and VHH-U (7KN5). The SARS-CoV-2 RBD is shown in a semi-transparent blue surface with the receptor-binding motif (RBM) highlighted in red. Other nanobodies are shown with colored lines as labeled.

**Figure 2.**
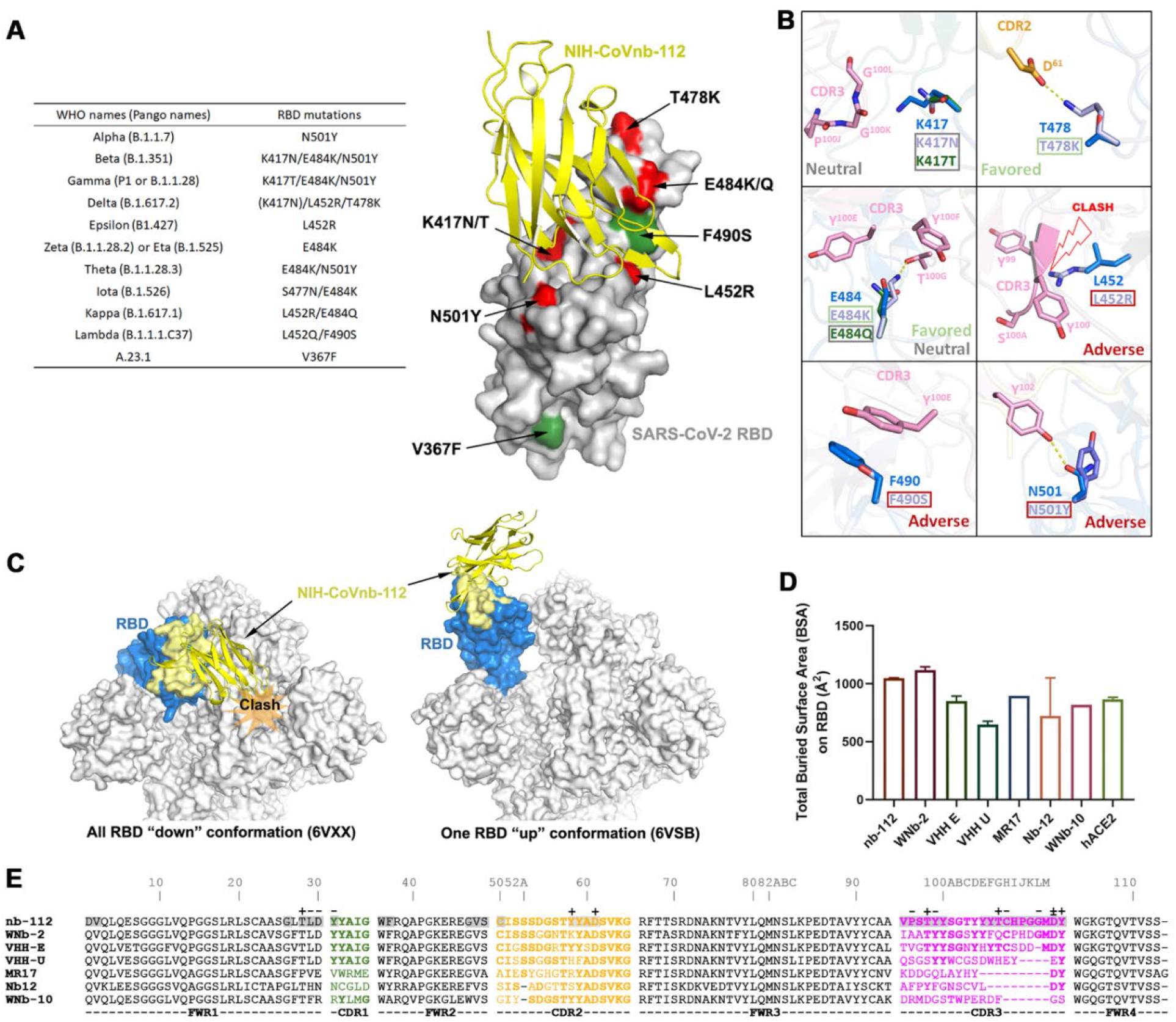
Crystal structure of NIH-CoVnb-112 in complex with SARS-CoV-2 RBD and effects of RBD mutations. (**a**) Diagram of RBD mutations occurred in the circulating SARS-CoV-2 variants. The recurrent RBD mutations within other SARS-CoV-2 variants are colored in red with unique mutations in green. (**b**) Composite model of RBD escape mutations on or around the epitope of NIH-CoVnb-112. The nanobody and prototype RBD are colored as in Fig. 1. The selected RBD mutations with side chain depicted only are shown in indicated colors. (**c**) RBD-based superimposition of the NIH-CoVnb-112-RBD complexes with SARS-CoV-2 spike in a closed (PDB: 6VXX)^46^ or one-RBD-up state (PDB: 6VSB).^47^ The nanobody is shown as yellow ribbons and the RBD domain is colored in blue with the nanobody epitope highlighted in yellow. The potential clashes of nanobody with the adjacent RBD of the closed spike trimer is shown in the left panel. (**d**) Comparisons between the RBD total buried surface areas (BSAs) of NIH-CoVnb-112, ACE2 and six published RBD-directed nanobodies. (**e**) Sequence alignments of NIH-CoVnb-112 with six structurally characterized RBD-binding nanobodies. The CDR sequences are colored as indicated in **Fig 1a**. The buried surface residues (BSA > 0 Å) as calculated by PISA are shaded in grey. Contact residues involved in salt-bridges or H-bonds to the RBD are marked above the sequence with (+) for the side chain and (−) for the main chain. The Kabat^19^ scheme was used to number nanobody amino acid residues, with unique insertion residues indicated by letter.

Outside of the CDR3’s extensive contact region, the extended CDR2 hairpin also helps stabilize the RBD tip (residues 481-487) (**Fig. 1c**-**d**). Of note, the side chain of F^486^_RBD_ stacks against the backbone of the CDR2 hairpin and fills the cavity formed by the inter-connected CDR2 and CDR3 loops. As a result, F^486^_RBD_ has the highest BSA value among all the RBD epitope residues (**Fig. 1d**). There are six additional hydrogen bonds formed between the RBD and nanobody residues outside of CDR3, including nanobody framework residue T^28^ to Q^498^_RBD_, CDR1 residues D^30^ and Y^31^ to G^445^_RBD_ and Y^449^_RBD_, and the side chains of CDR2 residues Y^58^ and D^61^ to Y^489^_RBD_ and N^487^_RBD_. Collectively, the CDR3-mediated nanobody-RBD recognition underlies the structural basis for NIH-CoVnb-112 neutralization, which overlaps the ACE2 binding site on the RBD.

The structure also provides information regarding NIH-CoVnb-112’s cross-reactivity against circulating SARS-CoV-2 escape variants. Based on the contribution of F^490^ to the interface we predict that the Lambda mutation F490S will disrupt π-π stacking with CDR3 Y^100E^ (**Fig. 2b**) which might lead to a minor impairment of NIH-CoVnb-112 recognition to this variant. On the other hand, the hydrogen-bond network between NIH-CoVnb-112 CDR3 residues and the RBD (residues 487-505) which ACE2 also uses in binding to the RBD bypasses the frequently mutated E^484^_RBD_ (**Fig. 1c**). The E484K or E484Q mutation present in the Beta, Gamma, Zeta, Theta, Iota and Kappa SARS-CoV-2 variants, can be accommodated by NIH-CoVnb-112 with no steric hindrance or charge-repulsion (**Fig. 2b**) which is supported by bio-layer interferometry (BLI) binding data showing that RBD_E484K_ binds to NIH-CoVnb-112 with a comparable affinity to that of wildtype (**Supplementary Fig. 1**). Another frequently mutated RBD residue, N^501^_RBD_, interacts with CDR3 Y^102^ with a hydrogen-bond distance of ∼2.9 Å. This can potentially be disrupted by the N501Y mutation present in the Alpha, Beta, Gamma and Theta SARS-CoV-2 variants (**Fig. 2a-b**), but BLI binding data shows only a 2-fold reduction in the K_D_ of RBD_N501Y_ as compared to wildtype (**Fig. 3b**). Notably, the K^417^_RBD_ mutation site also present in some of these variants is not involved in a hydrogen bond or salt bridge with NIH-CoVnb-112, suggesting that the K-to-N or K-to-T mutations here would have little impact on its binding (**Fig. 2b**). Interestingly, two Delta RBD mutations, L452R and T478K, may have opposite effects on NIH-CoVnb-112 binding: the extended guanidinium side chain of L452R likely clashes with CDR3 backbone while negatively charged T478K could reach out to the positively charged CDR2 D^61^ (**Fig. 2b**).

**Figure 3.**
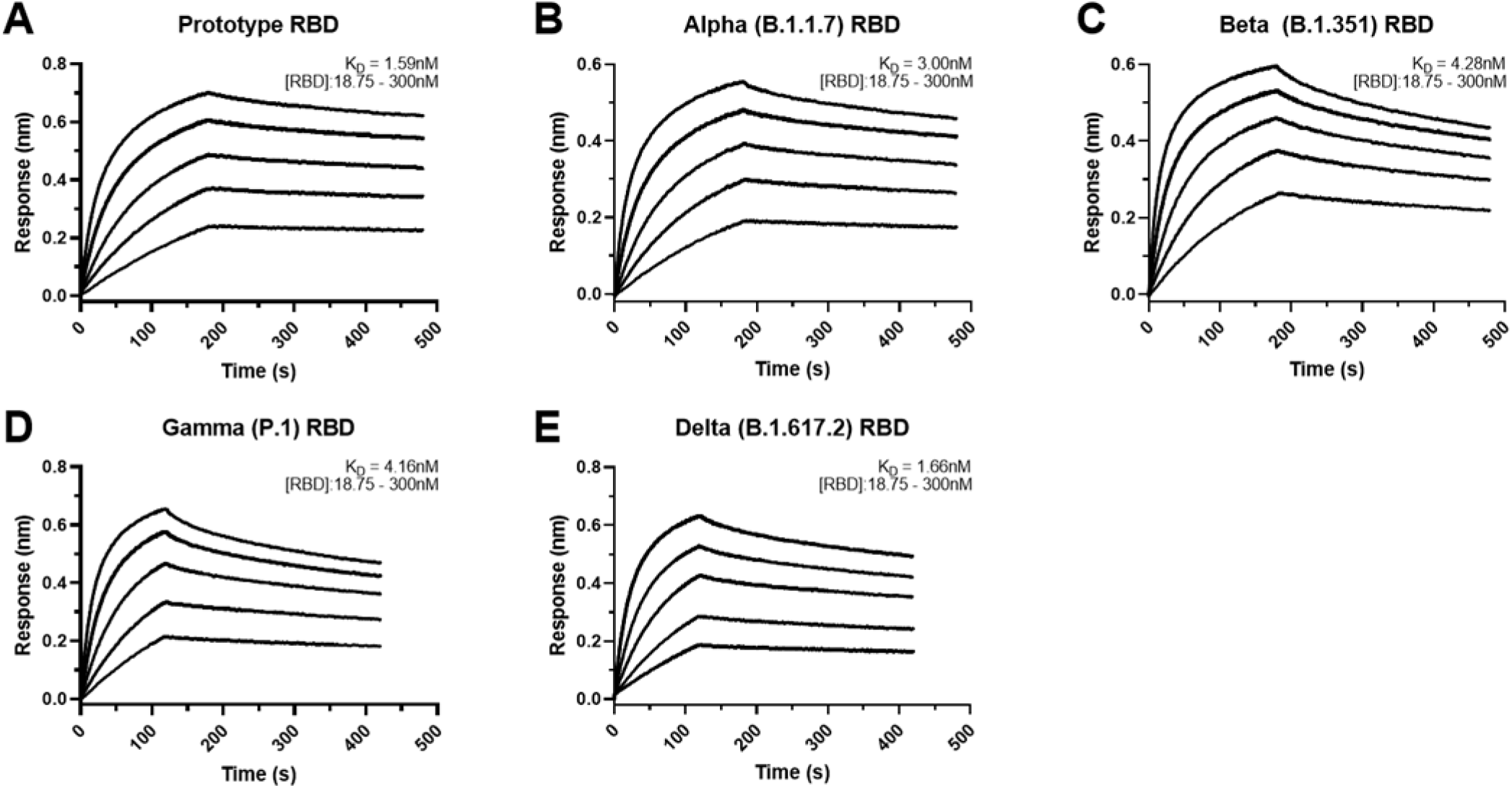
Affinity binding curves of NIH-CoVnb-112 against SARS-CoV-2 RBD variants. Using Biolayer Interferometry on a BioForte Octet Red96 system, association and dissociation rates were determined by immobilizing biotinylated-NIH-CoVnb-112 onto streptavidin-coated optical sensors (**a-e**). The nanobody-bound sensors were incubated with a concentration range (18.75 – 300nM) of recombinant RBDs, (**a**) Prototype, (**b**) Alpha (B.1.1.7), (**c**) Beta (B.1.351), (**d**) Gamma (P.1), and (**e**) Delta (B.1.617.2), for an initial time interval to allow association. The sensors were then moved to an RBD-free solution and allowed to dissociate over a second time interval. Curve fitting using a 1:1 interaction model allows for the affinity constant (K_D_) to be measured for each RBD variant as detailed in **Supplementary Table 2**.

The NIH-CoVnb-112 epitope largely overlaps with the binding sites of a number of other neutralizing antibodies (e.g., C105, CC12.1, CC12.3)^20^ and nanobodies (e.g., MR-17,^21^ WNb-2,^13^ and VHH-E^10^ **Fig. 1f**). Interestingly, two neutralizing nanobodies, alpaca-derived VHH-U^10^ and nanomouse-produced nb-12^11^, bind to the RBD core outside of the RBM and neutralize SARS-CoV-2 with a lower potency as compared with the best-in-class ACE2-blocking nanobodies suggesting that the RBM interaction is important for neutralization potency. Koenig et al. report that nanobody cocktails targeting distinct vulnerable epitopes on the RBD markedly suppress mutational escape^10^. Structural superposition of NIH-CoVnb-112 with the closed and open SARS-CoV-2 spike trimer reveals that it can only bind to the RBD in “up” conformation, even though its epitope remains mostly solvent accessible in the closed spike (**Fig. 2c**), which places it into the class 1 of RBD-specific antibodies/nanobodies.^20,22^ A cocktail of NIH-CoVnb-112 with nanobodies from other RBD binding classes might therefore suppress mutational escape. In total, there are 34 NIH-CoVnb-112 and 36 RBD residues involved in the binding interface with a buried surface area of ∼1045 Å^2^ for the complex, which is larger than those for other RBD specific nanobodies, yet similar in area to WNb-2, whose structures have been determined and significantly larger than that of the human receptor ACE2 for the RBD (**Fig. 2d**-**e**).

### *In vitro* affinity measurements of NIH-CoVnb-112 binding to SARS-CoV-2 RBD variants

To evaluate the potency of NIH-CoVnb-112 against the current variants of concern, we performed bio-layer interferometry using recombinant SARS-CoV-2 RBDs. NIH-CoVnb-112 was biotinylated using standard NHS-labeling chemistry to allow for capture on streptavidin-coated biosensors. Serial dilutions of each SARS-CoV-2 RBD, ranging from 300 – 18.75nM, were prepared in a sample buffer containing 0.1% BSA to block non-specific background binding. The background corrected curves were then fitted using a 1:1 interaction model to derive the equilibrium dissociation constants (K_D_), which results in a K_D_ measurement of 1.59nM (**Fig. 3a**). The affinities of several SARS-CoV-2 variant RBDs were then measured in the same manner with NIH-CoVnb-112 attached to sensor and variant RBDs in solution over the same concentration range. Curve fitting produced K_D_ values for Alpha: 3.0nM (**Fig. 3b**), Beta: 4.28nM (**Fig. 3c**), Gamma: 4.16nM (**Fig. 3d**), and Delta: 1.66nM (**Fig. 3e**). The individual association constants (k_on_) reveal similar values (**Supplementary Table 2**) of 1.53×10^5^ to 2.23×10^5^ M^- 1^s^-1^ for the prototype, Alpha, Beta, and Gamma RBDs, with the notable exception of the Delta variant RBD which had a 2-fold increase in the binding rate at 4.61×10^5^ M^-1^s^-1^. Conversely, all four variant RBDs had dissociation constants (k_off_) which were faster at 4.28×10^−4^ to 8.22×10^−4^ s^-1^, a 2 to 4-fold increase versus 2.36×10^−4^ s^-1^ for the prototype RBD. Thus, NIH-CoVnb-112 had similar *in vitro* binding to several relevant SARS-CoV-2 RBD variants.

### NIH-CoVnb-112 *in vitro* neutralization assessment of variant SARS-CoV-2 pseudovirus

To gain insight into the potential neutralization potency of NIH-CoVnb-112 against circulating SARS-CoV-2 variants of concern, we performed pseudotyped neutralization assays for several noted variants. Using the reported mutations and deletions for the variant spike proteins, modifications were introduced to the prototype sequence as follows: (Alpha - B.1.1.7 - Δ69-70, Δ144, N501Y, A570D, D614G, P681H, T716I, S982A and D1118H), (Beta - B.1.351 - D80A, Δ242-245, R246I, K417N, E484K, N501Y, D614G and A701V), (Gamma - P.1 - L18F, T20N, P26S, D138Y, R190S, K417T, E484K, N501Y, D614G, H655Y, T1027I and V1176F), and (Delta - B.1.617.2 - T19R, K77R, G142D, 156del, 157del, R158G, A222V, L452R, T478K, D614G, P681R and D950N). Pseudotyped lentivirus possessing an RFP reporter was produced by transient transfection of HEK293T/17 cells and the virons were harvested and purified over a sucrose cushion. For each variant, a multiplicity of infection of 0.5 was used to transduce monolayers of HEK293T-ACE2 cells in the presence of NIH-CoVnb-112 at concentrations ranging from 714pM to 714nM. Media was replaced at 24 hours and the cells harvested at 48 hours post infection for flow cytometry. The percentage of transduction, as measured by the gated positive RFP population, for each concentration was normalized using a virus only transduction at the time of each assay to calculate percent inhibition. Dose-response curve fitting (**Fig. 4**) allows for calculating the half-maximal effective concentration (EC_50_) for each variant pseudovirus. The EC_50_ for the SARS-CoV-2 prototype sequence (14.3nM) is within agreement of the previously reported concentration.^9^ The SARS-CoV-2 variant sequences similarly yielded EC_50_ values: Alpha (9.4nM), Beta (15.8nM), Gamma (17.6nM) and Delta (14.5nM). Notably, there are differences in the apparent slopes (**Fig. 4** and **Supplementary Table 3**) of the neutralization curves, which have been correlated with the cooperativity of binding.^23^ Thus, NIH-CoVnb-112 had similar potency in pseudotyped neutralization assays for several relevant SARS-CoV-2 spike variants.

**Figure 4.**
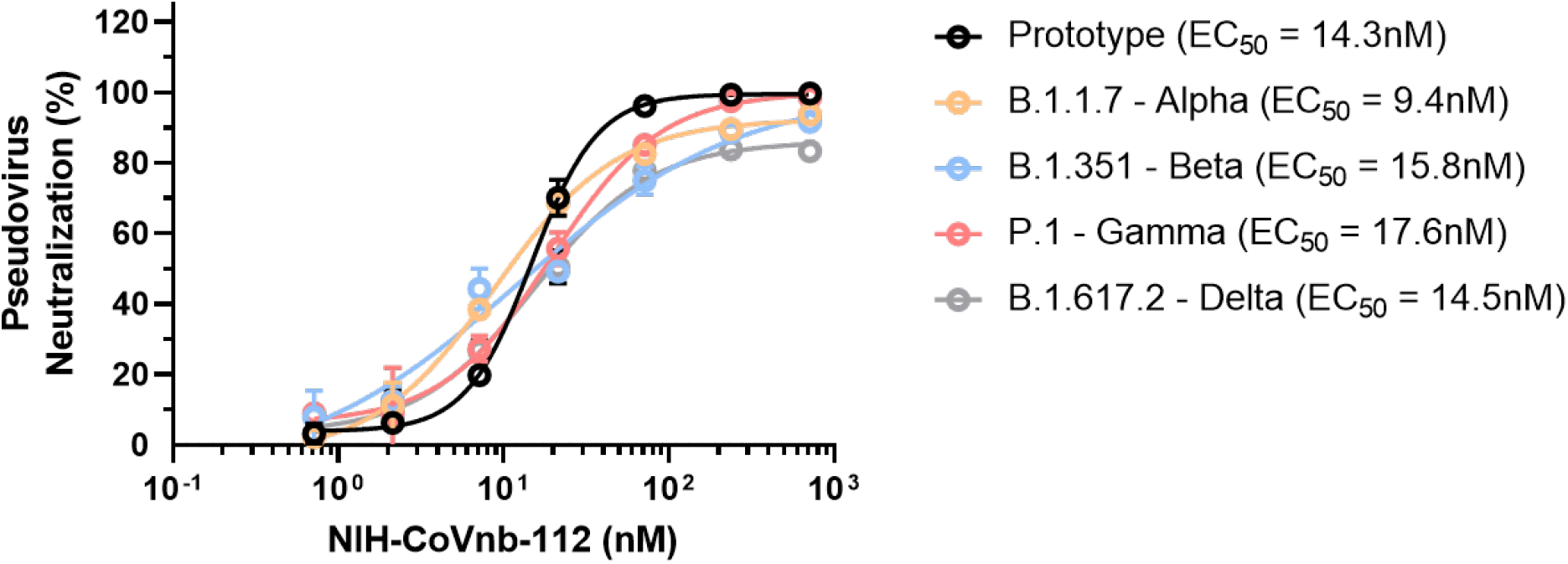
NIH-CoVnb-112 inhibition of SARS-CoV-2 variant pseudovirus following nebulization. HEK293 cells overexpressing human ACE2 were cultured for 24 hours with pseudotyped SARS-CoV-2 spike variant virus, pretreated with NIH-CoVnb-112 at different concentrations. Following 48 hours incubation, HEK293-ACE2 cells were analyzed by flow cytometry to quantify the fluorescence level. NIH-CoVnb-112 potently inhibited viral transduction for all variant spike pseudoviruses with the following EC50 values: Prototype (14.2nM), Alpha (9.4nM), Beta (15.8nM), Gamma (17.6nM), and Delta (14.5nM), respectively. Curve fit parameters are detailed in **Supplementary Table 3**.

### Nebulization treatment of NIH-CoVnb-112 in a hamster model of COVID-19

To assess the potential prophylactic and therapeutic efficacy of nebulized nanobody administration, we chose to evaluate nanobody prophylaxis (−24 hour pre-infection) and multiple post-exposure (+12 hour., 1dpi, 2dpi) treatments as outlined (**Fig. 5a**). Hamsters were randomized into saline only (n=8) or NIH-CoVnb-112 (n=8) only. Twenty-four hours prior to virus challenge, groups of 4 animals were placed in a containment box with an attached nebulizer. Animals received nebulized exposure of a 5mL solution containing normal saline alone or normal saline with 25mg/mL NIH-CoVnb-112 for 20 minutes. Twenty-four hours after the initial dose, all animals were challenged with ∼10^4^ pfu of prototype SARS-CoV-2 by intranasal administration. Additional nebulization doses were given at 12 hours post virus infection, at 1 day post infection and for half of each group an additional dose at 2 days post infection as indicated (**Fig. 5a**). This design was selected based on an initial pilot study (**Supplementary Figs. 3-7**). Animals were observed for weight on days 1, 2, 3, 5, and 7 post infection (**Fig. 5b**) and oropharynx swabs taken on days 1, 2, and 3 post infection (**Supplementary Fig. 7**). Half of each group was euthanized on day 3 post infection and the remaining half euthanized on day 7 post infection for assessments of viral titer and histopathology assessments. Baseline weights of the nanobody group (169.3, 171.6, 192.6, and 164.9 grams) and the saline group (185.0, 199.7, 171.2, and 186.2 grams) were recorded prior to nebulization and compared to day 7, nanobody group (169.6, 168.5, 196.7, and 164.8 grams) and the saline group (168.2, 179.2, 162.2, and 175.3 grams), reveal an initial reduction for both groups (**Fig. 5b**) followed by resolution to near baseline in the NIH-CoVnb-112 treated group. In both groups, nasal turbinates at 3 days post infection (**Fig. 5c**) had a sustained level of viral burden, with no apparent difference between treatment groups. By contrast, the level of viral burden in the cranial lung (**Fig. 5d**) had a striking decrease of 6 order of magnitude, with three of the four samples below the limit of detection of the plaque assay (LLOQ = 10 pfu/100mg). In the oropharyngeal swabs (**Supplementary Fig. 7**), both groups follow a decrease in viral titer over time with no apparent difference between groups. To assess the delivery dose to the lower respiratory tract following nebulization, hamsters euthanized on day 3 post infection had bronchoalveolar lavage performed and ELISA used to measure NIH-CoVnb-112 concentration. The level of NIH-CoVnb-112 on day 3 post infection (**Supplementary Fig. 8**) had a mean concentration of 154 ng/mL, while the saline group, as expected, yielded only background values below the lower limit of quantification. Lung and tracheal tissue were collected at days 3 and 7 post infection to assess the impact of nebulization delivery of NIH-CoVnb-112 on pathology as described above using the semi-quantitative lung scoring scale on H&E sections in a blinded manner (**Supplementary Table 5, Fig. 5e-f, and Supplementary Fig. 9**). Histological observation of the saline treatment group revealed pathological evidence of severe pulmonary disease, including perivascular and peribronchial mononuclear cell infiltration, focal tracheal necrosis and exfoliation, and moderate to severe septal and intra-alveolar infiltration of mononuclear cells (**Fig. 6**). In addition, marked interstitial inflammation and pneumocyte hyperplasia were prominent in the saline treatment group (**Supplementary Fig. 10**). In the NIH-CoVnb-112 treatment group, at both days 3 and 7 post infection the histology was within normal limits, with minimal tracheal and perivascular mononuclear cell infiltration and accumulation on day 7 (**Fig. 6**). The total lung score and subset scores (**Fig. 5e-f and Supplementary Fig. 9**) reveal a pronounced distinction in lung score for the NIH-CoVnb-112 group (day 3 mean score = 2) versus the saline group (day 3 mean score = 26.75). By day 7 there was a modest increase in the NIH-CoVnb-112 group (day 7 mean score = 7), while the saline group increased markedly (day 7 mean score = 39.75). The increase in NIH-CoVnb-112 treatment lung score was a result of minor increases in vascular, alveolar, and interstitial inflammation (**Supplementary Fig. 9**). Thus, nebulized NIH-CoVnb-112 demonstrated initial *in vivo* protective efficacy in a hamster model of SARS-CoV-2 infection.

**Figure 5.**
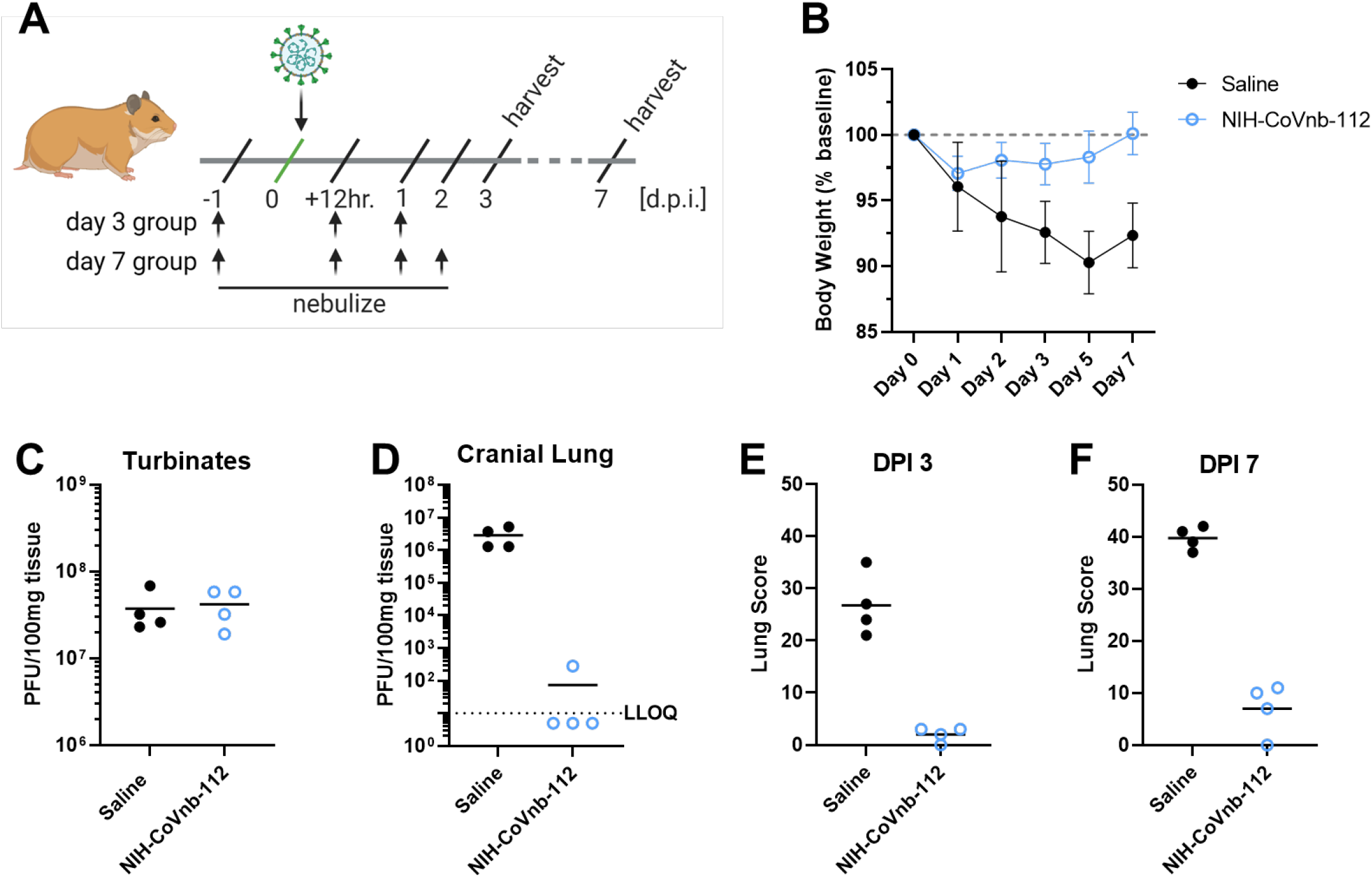
NIH-CoVnb-112 nebulization treatment in a hamster model of SARS-CoV-2. (**a**) Schematic overview of study design for nebulization treatment of Syrian hamsters with NIH-CoVnb-112. Adult Syrian hamsters (n=8/group, males, 12 weeks old) were treated at -24 hours prior to virus challenge with nebulized saline or NIH-CoVnb-112 (25mg/mL) in a 5mL volume over 20 minutes. Following intranasal challenge, each group was treated at 12 hours post infection with the same condition, and again at 1 dpi. Half of the animals in each group received a fourth dose at 2 dpi. Animals were weighed daily and oral swabs taken at dpi 1-3. Groups were euthanized at day 3 and day 7 post infection respectively and sample taken for assessment. (Figure elements generated using BioRender.com) (**b**) Body weight change as a percentage of baseline weight for day 7 post infection group (n=4/group, mean +/- SD) (**c**) Viral burden from turbinate tissue of the day 7 post infection group as determined by double plaque overlay assay. (**d**) Viral burden from cranial lung tissue of the day 7 post infection group as determined by double plaque overlay assay. (**e-f**) Hematoxylin and eosin-stained tissue sections from the day 3 and day 7 post infection animals were semi-quantitively scored by a pathologist blinded to condition using an established scoring metric. Metrics included overall lesion extent, bronchitis, alveolitis, pneumocyte hyperplasia, vasculitis, and interstitial inflammation; each on a 0–4 or 0–5 scale and scores summed. Detailed data for day 3 and day 7 post infection are shown in **Supplementary Fig. 9**.

**Figure 6.**
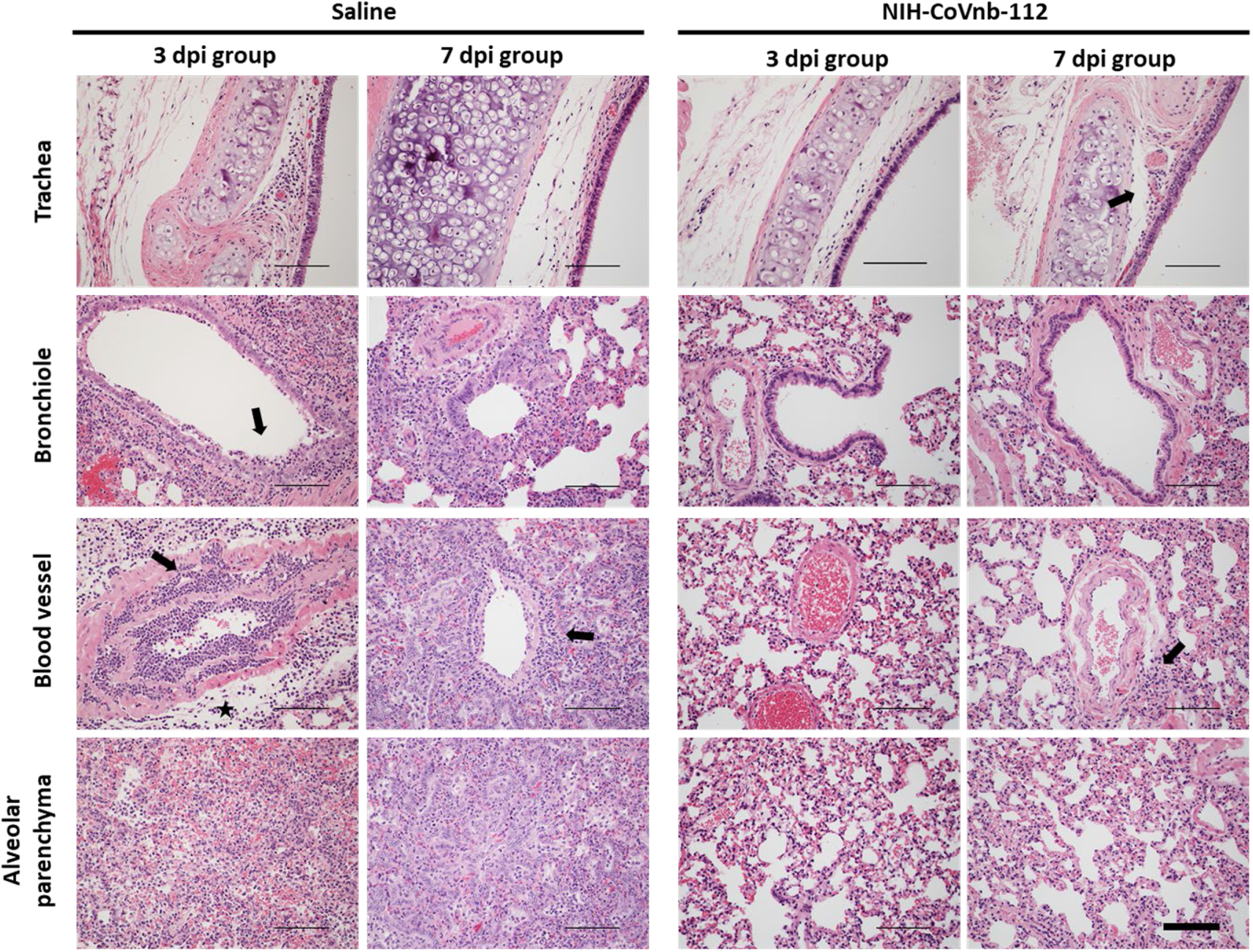
Histopathological staining of trachea and lung sections from multi dose nebulization treated Syrian hamsters following SARS-CoV-2 infection. Representative hematoxylin and eosin-stained tissue sections from day 3 and day 7 post infection groups (n = 4 per group) for trachea, bronchioles, blood vessels, and alveolar parenchyma. Pathology of note for the saline group includes epithelial cell exfoliation and cellular debris accumulation in the bronchiole lumen (3dpi, arrow), mononuclear cell and heterophile infiltration and accumulation of leukocytes in blood vessel smooth muscle layers (3dpi, arrow), severe perivascular edema (3dpi, star) and transmural leukocyte migration and perivascular accumulation (7dpi, arrow). The nanobody treated group pathology remained within normal limits at day 3 post infection, exhibiting minor pathology of note on day 7 including minimal mononuclear cell infiltration in the tracheal subepithelial interstitium (7dpi, arrow) and minimal perivascular accumulation of mononuclear cells (7dpi, arrow). Corresponding pathologist descriptions found in **Supplementary Table 5**. Representative images for each group column are from the same animal. Images acquired at 200x magnification. Scale bars for all images = 300µm.

## DISCUSSION

In summary, we provide the structural basis for effective neutralization of the SARS-CoV-2 virus and current variants of concern using the previously isolated nanobody, NIH-CoVnb-112.^9^ We determined the crystal structure of NIH-CoVnb-112 complexed with its primary viral target, the SARS-CoV-2 RBD, to 2.8 Å resolution. We further explored the neutralization potential by performing pseudovirus neutralization assays with lentivirus expressing spike proteins with the entire constellation of mutations reported for each variant. We observed robust neutralization of each variant in a cell culture transduction assay, with EC50 values comparable to the prototype SARS-CoV-2 spike lentivirus ranging from 9.4 – 17.6 nM. We then performed initial studies to determine the ability of the monomeric version of NIH-CoVnb-112 to provide protection when delivered by nebulization prior to and following viral challenge in a Syrian hamster model of SARS-CoV-2 infection. We observed six orders of magnitude reduction in the viral lung burden, absence of gross weight loss, and a significant reduction of lung pathology, demonstrating a robust protective benefit of nebulized nanobody therapy.

Given their small size and robust thermostability, nanobodies can be aerosolized for the direct treatment of respiratory diseases, as in the case of delivery of a trimeric nanobody for the treatment of respiratory syncytial virus.^24–26^ Previous work has established the hamster as a useful model of COVID-19 disease, owing to robust viral replication in the respiratory tract, which results in weight loss and significant lung pathology in a reliable and consistent manner.^27^ Recent effort has demonstrated the ability of nanobodies in various formats to effectively inhibit the SARS-CoV-2 virus in both prophylactic and therapeutic settings in rodent models of COVID-19.^10,11,26,28–31^ Most studies have utilized nanobody multimers or nanobody Fc-fusions for intraperitoneal or intranasal instillation delivery, which have demonstrated effective protection against lung pathology and viral burden in the respiratory tract.^28–30^These studies did not demonstrate therapeutic delivery via nebulization. In contrast, Nambulli *et al*. delivered a trimeric nanobody, termed Pin-21, by nebulization in a hamster model of COVID-19 and observed a robust reduction in both lung pathology and viral burden with the maintenance of body weight during the study. These studies and our work provide support for the further investigation of nanobody-based pulmonary therapeutics for SARS-CoV-2 infection.

We envision the use of nebulized nanobody-based treatments to provide prophylactic protection in healthcare settings, such as long-term care facilities, potentially reducing viral transmission in these community settings. In addition, the use of nebulized nanobody therapeutics could hypothetically provide effective early phase treatment in SARS-CoV-2 infections, delivered in the home setting, potentially reducing the severity of disease. Translation to clinical trials will require improved delivery in animal models, including assessment in non-human primates, which represent a closer approximation to human lung anatomy. In addition, it is imperative to explore potential toxicity and immunogenicity effects due to nebulized delivery to ensure a satisfactory safety profile prior to human clinical trials.

### Limitations of the study

While the current results provide strength for using nanobodies as potential inhaled therapeutics, there are several limitations to this current study. First, the use of bio-layer interferometry and pseudovirus assays to measure *in vitro* binding and neutralization provide evidence that NIH-CoVnb-112 has broad potency, yet these assays must be interpreted with caution. In the setting of authentic SARS-CoV-2 infection, and particularly the highly transmissible Delta variant, it is vital to assess the actual virus to ensure parity between *in vitro* and *in vivo* findings. The constellation of mutations in variants outside of the RBD may provide competitive fitness, which outstrips the ability of NIH-CoVnb-112 to effectively neutralize infection. Second, while acute weight loss and lung pathology provide metrics for assessing the early phase of infection, the natural course of disease in the Syrian hamster resolves with rare mortality.^27^ Thus, the hamster model should be considered just one of several relevant *in vivo* models. Third, the hamster experiments should be extended for future studies; the experiments presented here serve primarily as proof-of-concept for prophylactic and therapeutic potential. Given resource limitations, our studies had limitations on sample size and thus we have reported our findings without statistical analysis. We observed a robust viral load reduction and clear impact on lung pathology, which is encouraging but should not be considered definitive. Fourth, a purely therapeutic, multidose group with nebulization starting after symptomatic infection was not performed in this study and should be considered for future experiments to assess post-infection therapeutic efficacy. Fifth, dose-response studies would help inform the effective dose range of the delivered nanobody. Sixth, we did not assess NIH-CoVnb-112 clearance from the lung into the blood or the potential immunogenicity response from multiple dosing. Seventh, nebulizing NIH-CoVnb-112 in a passive containment system likely is inefficient for delivery to the upper and lower respiratory airways. Indeed, much of the nebulized nanobody was observed to condense onto the walls of the containment box and the fur of the hamsters. Use of a directed inhalation device would decrease the amount of nanobody used and provide a standard dose delivered. Eighth, the monomeric nature of NIH-CoVnb-112 may lead to reduced residence time in the lung, therefore methods for increasing the half-life should be explored to reduce the concentration required and number of doses for therapeutic effect. Exploring a multivalent nanobody cocktail^12,13^ of non-overlapping epitope nanobodies or multimerized^10,11,32,33^ formatting of the lead nanobody may increase efficiency as reported by others and reduce the risk of immune escape. Other approaches such as *in vitro* affinity maturation^16,34,35^ may be employed to significantly improve on the existing affinity.

In conclusion, we have demonstrated that NIH-CoVnb-112 possesses broad binding and neutralization capacity in the context of the current circulating SARS-CoV-2 variants. These results provide support for continued development of NIH-CoVnb-112 as a nebulized delivery adjunct therapeutic for reducing early phase viral burden and prophylaxis against lung pathology.

## MATERIALS AND METHODS

### SARS-CoV-2 RBD production

The expression plasmid encoding for codon-optimized SARS-CoV-2 RBD (329-538) with a C-terminal hexa-histidine tag was transfected into Expi293™ GnTI-(Thermo Fisher) cells (1×10^6^ cells/mL) with PEI-Max. One week post-transfection, the clarified supernatant was purified on Ni-NTA column (Cytiva), followed by size-exclusion chromatography on HiLoad 26/600 Superdex 200 (Cytiva) equilibrated with PBS buffered saline pH 7.4.

### Crystallography and structure determination of NIH-CoVnb-112 and SARS-CoV-2 RBD

The purified SARS-CoV-2 RBD (329-538) was mixed with an excess of NIH-CoVnb-112 (molar ratio 1:4) and incubated at 4° C overnight before separation on HiLoad 26/600 Superdex 200 column (Cytiva) which was pre-equilibrated in 10mM Tris pH 8.0 and 100mM ammonium acetate. The complex fractions were pooled and concentrated to ∼13 mg/mL for crystallization. Crystallization trials of the nanbody-RBD complex were performed using the vapor-diffusion hanging drop method with the sparse matrix crystallization screens ProPlex (Molecular Dimensions), Index (Hampton Research), and Crystal Screen I and II (Hampton Research) with a 1:1 ratio of protein to well solution. After approximately 3 weeks incubation at 21°C, plate-shaped crystals were obtained in 0.1 M magnesium acetate, 0.1 M MOPS pH 7.5, 12% PEG 8000. Crystals were snap-frozen in the crystallization condition supplemented with 20% 2-methyl-2, 4-pentanediol (MPD) used as the cryoprotectant. X-ray diffraction data was collected at the SSRL beamline 9-2 and was processed with HKL3000.^36^ The structure was solved by molecular replacement in PHASER from the CCP4 suite^37^ using 7LDJ as a template in which the RBD moiety and nanobody were treated as independent searching models. Iterative cycles of model building and refinement were done in Coot^38^ and Phenix.^39^ Data collection and refinement statistics are provided in **Supplementary Table 1**. Structural analysis and figure generation were performed in PyMOL.

### Structure validation and analysis

Ramachandran statistics were calculated with MolProbity^40^ and illustrations were prepared with PyMOL Molecular graphics (http://pymol.org). Nanobody-RBD interface and buried surface area were determined in PISA.^41^ The complex structure was deposited in the Protein Data Bank with accession code 7RBY.

### Scaled fermentation expression of NIH-CoVnb-112 in *Pichai pastoris*

Using the previously characterized^9^ strain of X-33 *Pichia pastoris* expressing NIH-CoVnb-112, fermentation was performed to produce sufficient material for in vivo studies and reduce the risk of endotoxin contamination. Fermentation was performed by the University of Georgia Bioexpression and Fermentation Facility (UGA BFF, Athens, GA) and performed as briefly follows: A 5-10% inoculum of the expression clone in buffered glycerol complex medium (BMGY) was grown for 24 hours at 30°C. The overnight culture (300mL) was used to inoculate a 4-L DCI Tryton bioreactor containing 2.7L basal salts medium (BSM) for a 75% working volume. Post-batch growth was fed with 50% v/v glycerol until the wet cell weight reached 200grams/L at which point feed was switched to methanol containing 12mL/L *Pichia* trace metal salts for induction of expression. Final methanol concentration was maintained in the fermenter at 1gram/L for 72 hours. Following harvest and clarification, the supernatant volume was reduced 10-fold and exchanged to buffer (20mMTris, pH 7.4, 0.5M NaCl) using tangential flow filtration. NIH-CoVnb-112 was affinity purified using a 5mL HiTrap HP column (Cytiva 17115201) charged with nickel. The sample was eluted using a linear gradient of imidazole to 0.5M. The eluted nanobody was buffer exchanged to 0.9% normal saline. and the final concentration determined based on ProtParam^42^ mass extinction coefficient estimate (1 mg/mL solution will have an A280 absorbance value of 2.0310) for a yield of 1.1g. The purity (95.5%) was determined using SDS-PAGE. Endotoxin level was measured using the LAL Chromogenic Endotoxin Quantitation Kit (Cat# 88282, Thermo Scientific) to verify a concentration of less than 0.1 endotoxin unit/mL (EU/mL) of purified nanobody.

### Pseudovirus Neutralization Assay of SARS-CoV-2 Variants of Concern

Neutralization potency of NIH-CoVnb-112 was assessed using the method previously reported^9^ with modifications to the SARS-CoV-2 spike protein sequence. Briefly, lentiviruses were propagated in HEK293T/17 cells (ATCC # CRL-11268) following published Current Protocols in Neuroscience.^43^ Plasmids for SARS-CoV-2 variants were synthesized using the prototype sequence (GenScript MC_0101081, human codon optimized, ER retention signal removed) as the reference sequence. SARS-CoV-2 variants included the following mutations: (Alpha - B.1.1.7 - Δ69-70, Δ144, N501Y, A570D, D614G, P681H, T716I, S982A and D1118H), (Beta - B.1.351 - D80A, Δ242-245, R246I, K417N, E484K, N501Y, D614G and A701V), (Gamma - P.1 - L18F, T20N, P26S, D138Y, R190S, K417T, E484K, N501Y, D614G, H655Y, T1027I and V1176F), and (Delta - B.1.617.2 - T19R, K77R, G142D, 156del, 157del, R158G, A222V, L452R, T478K, D614G, P681R and D950N). Monolayer cultures of 293T cells were transiently transfected with plasmids expressing SARS-CoV-2 spike protein variants, psPAX2 (Addgene #12260), and a lentiviral transfer vector CD512-EF1a-RFP (System Biosciences CD512B-1) using Lipofectamine 2000. Supernatant was harvested 48 hours post transfection and concentrated by centrifugation at 50,000 g for 2 hours over a 20% sucrose cushion. Pellets were resuspended in PBS and used for infection. Titers were determined by performing biological titration of fluorescent viruses by flow cytometry to ensure uniformity of assays.

For transduction assays, HEK293Ts expressing human angiotensin-converting enzyme 2 (HEK293T- ACE2, BEI Resources, NR-52511) were plated at the density of 50,000 cells/well in a 24-well plate. Cells were transduced with SARS-CoV-2 pseudotyped recombinant lentiviruses expressing RFP (S-CD512-EF1a-RFP) with a multiplicity of infection (MOI) of 0.5 +/- NIH-CoVnb-112 at described concentrations. Media on cells was replaced the next day. 48 hours post transduction, cells were trypsinized and fixed in 1% formaldehyde. BD LSRFortess Flow Cytometer was used to measure percent fluorescent cells and the mean fluorescent intensity per replicate. The assays were performed blinded using the diluent as a negative control. All experiments were performed in triplicate.

### Bio-layer Interferometry affinity measurement of NIH-CoVnb-112 binding variant SARS-CoV-2 RBDs

Bio-layer interferometry was used to measure the affinity binding constants of *Pichia pastoris* expressed NIH-CoVnb-112 against SARS-CoV-2 variant RBD proteins. NIH-CoVnb-112 was biotinylated using NHS-EZ-Link Sulfo-NHS-LC-Biotin (#21335, ThermoFisher) with a degree of labeling of 1 to reduce impact on binding. Assay conditions were prepared using 0.1% BSA (w/v) in 1xPBS as the assay and blocking buffer in a volume of 300 µL. Biotinylated-NIH-CoVnb-112 was diluted into assay buffer at 5 µg/mL and immobilized onto streptavidin coated biosensors (#18-5019, Sartorius) to a minimum response value of 1 nm on the Octet Red96 System (Sartorius). A baseline response was established in assay buffer prior to each association. Recombinant SARS-CoV-2 RBDs [B.1.1.7/N501Y (SPD-C52Hn, ACRO Biosystems); B.1.351/K417N, E484K, N501Y (SPD-C52Hp, ACRO Biosystems); P.1/K417T, E484K, N501Y (SPD-C52Hr, ACRO Biosystems); B.1.617.2/L452R, T478K (SPD-C52Hh, ACRO Biosystems); L452R (SPD-C52He, ACRO Biosystems); K417N (SPD-C52Hs, ACRO Biosystems); E484K (SRD-C52H3, ACRO Biosystems); T478K (40592-V08H91, Sino Biological)] were diluted into assay buffer at the specified concentrations. The variant SARS-CoV-2 RBDs were associated for 120s followed by dissociation for 300s in the same baseline wells. The assay included one biosensor with only assay buffer which was used as the background normalization control. Using the ForteBio Data Analysis suite, the data was normalized to the association curves following background normalization and Savitzky-Golay filtering. Curve fitting was applied using global fitting of the sensor data and a steady state analysis calculated to determine the association and dissociation constants as reported in **Supplementary Table 2**.

### NIH-CoVnb-112 Post-nebulization stability evaluation

Stability following nebulization of *Pichia pastoris* expressed NIH-CoVnb-112 was performed using an PARI Sprint nebulizer placed in line with a custom glass bead condenser. As previously described,^9^ a plastic culture tube was fitted with a glass-pore frit and filled with sterilized 5mm borosilicate glass beads. A three-way stopcock was positioned distal to the frit to prevent pressurization during sample nebulization. A 2mg/mL NIH-CoVnb-112 solution was prepared in 0.9% normal and nebulized into the condenser. The nebulized nanobody was collected and protein concentration measured by A280 absorbance. Pre- and post-nebulization samples were injected onto a Superdex 75 Increase 10/300 GL size exclusion column (SEC) operating on an AKTA Pure 25 M system to assess stability and aggregation propensity.

### Animal use ethics statement

Hamsters were used according to protocols approved by the Institutional Animal Care and Use Committees of Colorado State University (CSU).

### Animal experimental procedures

Male Syrian hamsters (*Mesocricetus auratus*) were purchased from Charles River Laboratories and used at 12 weeks of age. Hamsters were moved to ABSL3 6–8 days prior to challenge. All animals were challenged with ∼10^4^ pfu of SARS-CoV-2 (2019-nCoV/USA-WA1/2020 strain) under light anesthesia with ketamine-xylazine. Virus diluted in PBS was administered via pipette into the nares (100 µl total, ∼50 µl/nare); animals were observed until fully recovered from anesthesia. Virus back-titration was performed on Vero E6 cells immediately following inoculation, confirming that hamsters received 9.4 × 10^3^ pfu. Animals (*n* = 6-8/group) were monitored daily post infection for clinical signs of infection (weight loss, nasal discharge, etc.) and weighed daily until euthanasia. Oropharyngeal swabs were collected from all hamsters on days 1, 2, and 3 post infection and assayed for infectious virus titer by plaque assay. Half the animals (3-4 hamsters) in each group were euthanized at Day 3 post infection (acute phase) to assess nasal turbinate and lung (cranial lobe) virus titer and histopathology, and the other half of each group euthanized at Day 7 (subacute phase) post infection to assess histopathology.

### NIH-CoVnb-112 nebulization in hamsters

Aerosol delivery of NIH-CoVb-112 or normal saline was performed using a PARI LC Sprint nebulizer attached to a housing unit approximately 30×18×10cm with fitted out-flow filter. Groups of 3-4 animals were dosed for 15-20 minutes with a 5mL volume containing normal saline or nanobody solution. An initial cohort of animals (n = 6/group; 5 groups) were dosed 24 hours prior to virus challenge and 8 hours post infection with either normal saline or NIH-CoVnb-112 solution (10mg/mL) as follows: saline/saline, saline/nanobody, nanobody/saline, or nanobody/nanobody. A control group receiving no nebulization exposure was included. The sample size was based on the number of available animals and resources of the collaborating investigator and not determined by a formal power calculation. The animals were monitored, and samples collected as described above. A second cohort of animals (n = 8/group; 2 groups) were dosed 24 hours prior to virus infection with either normal saline or NIH-CoVnb-112 solution (25mg/mL). Repeat dosing was performed using the same solution at 12 hours post virus infection, and 1dpi. Half of the animals (n = 4/group) per group received an additional nebulization dose at 2dpi. The animals were monitored, and samples collected as described above. In addition, bronchoalveolar lavage fluid (BALF) was collected on the day 3 post infection group to assess NIH-CoVnb-112 concentration resident in the lower respiratory tract. All nebulization procedures were performed in the ABSL3 facility. Samples were double blinded, with an independent investigator blinding the containers holding nanobody or saline treatments and hamster samples prior to transfer to collaborators and holding the sample key until the completion of all analysis. All hamsters were included in the study and analysis; no animals died or were excluded from data analysis.

### Quantification of SARS-CoV-2 in procedure samples

Virus titration was performed on oropharyngeal swabs obtained at 1, 2, and 3 days post infection and on cranial lung tissue samples obtained at 7 days post infection by double-overlay plaque assay on Vero E6 cells as previously described.^44^ Briefly, fluids or tissue homogenates were serially diluted in Tris-buffered Minimum Essential Medium (MEM) with 1% BSA and inoculated onto confluent monolayers of Vero E6 cells seeded in 6-well cell culture plates; incubated at 37 °C for 45 min; and each well overlaid with 2 ml of MEM containing 2% fetal bovine serum and 0.5% agarose. After 24–30 hour incubation at 37 °C in 5% CO_2_, a second overlay containing neutral red dye was added, and plaques were counted 24-36 hours later with the aid of a lightbox.

### Measurement of NIH-CoVnb-112 in bronchoalveolar lavage fluid

SARS-CoV-2 RBD (#SPD-C52H3, ACRO Biosystems) was coated at 2 µg/mL in 1xPhosphate Buffered Saline, 100 µL per well, onto Nunc Maxisorp plates overnight at 4°C. Coating solution was removed, and plate blocked with 300 µL 4% non-fat dry milk in 1xPBS for 1 hour at room temperature. Serial dilutions of BALF and NIH-CoVnb-112 as the standard (1.95 – 125ng/mL) were prepared in 1% non-fat dry milk in 1xPBS and transferred to the blocked plate in triplicate. The samples were incubated for 1 hour to allow for association with RBD. The assay plate was washed and 100 µL peroxidase conjugated goat anti-alpaca VHH domain-specific antibody (#128-035-232, Jackson ImmunoResearch), diluted to 0.8 µg/mL, transferred into each well and incubated for 1 hour at room temperature. After a final wash, the plate was developed by the addition of 100 µL tetramethylbenzidine (TMB) (#T5569, Sigma-Aldrich). Assay development was stopped by addition of 50µL 1M sulfuric acid and the assay absorbance measured at 450 nm on a Biotek Synergy 2 plate reader. The assay plate was washed with 1xPBS five times between each step of the assay.

### Histopathological assessment and quantitative scoring of lung pathology

Tissues from hamsters were fixed in 10% buffered formalin for 7–14 days, embedded in paraffin, and cut sections stained with hematoxylin and eosin (H&E). Slides were read by a single veterinary pathologist blinded to the identity of the group including both day post infection and treatment status. Cranial lung histopathology was scored as previously described^45^ for the following metrics: overall lesion extent, bronchitis, alveolitis, pneumocyte hyperplasia, vasculitis, and interstitial inflammation; each on a 0–4 or 0–5 scale, and scores summed for each animal. Repeated intra-rater reliability scoring assessments deviated less than 5% of reported values.

## ACKNOWLEDGEMENT

Use of the Stanford Synchrotron Radiation Lightsource, SLAC National Accelerator Laboratory, is supported by the U.S. Department of Energy, Office of Science, Office of Basic Energy Sciences under Contract No. DE-AC02-76SF00515. The SSRL Structural Molecular Biology Program is supported by the DOE Office of Biological and Environmental Research, and by the National Institutes of Health, National Institute of General Medical Sciences. The study was supported by the NINDS intramural research program under the Laboratory of Functional and Molecular Imaging, directed by Dr. Alan Koretsky. The funders had no role in study design, data collection and analysis, decision to publish, or preparation of the manuscript and the contents of this publication are solely the responsibility of the authors. We thank the NHLBI Biophysics Core Facility for the use of BLI instrument. We thank Dr. Thomas Stanley in the NIEHS Structural Biology Core for construction of envelope plasmids. We thank Dr. Yoshima Akahata for assistance with animal study blinding. A provisional patent covering NIH-CoVnb-112 and associated nanobody sequences was filed (U.S. Provisional Application No.: 63/055,865, Filing Date July 23, 2020) with a PCT patent application (application number PCT/US21/42883) filed on July 23, 2021.

## AUTHOR CONTRIBUTIONS STATEMENT

DLB, TJE, RAB, and MP conceived the experiments. TJE, YZC, NPM, WDT, RAB, performed experiments and analysis. HBO provided blinded histopathology analysis. DLB, TJE, MP, and YZC wrote the manuscript.

## DISCLAIMER

The opinions and assertions expressed herein are those of the authors and do not necessarily reflect the official policy or position of the Uniformed Services University (USU), the Department of Defense (DoD), the National Institutes of Health (NIH) or any other US government agency, or the Henry M. Jackson Foundation for the Advancement of Military Medicine, Inc.

## REFERENCES

1 Zhou, D. et al. Robust SARS-CoV-2 infection in nasal turbinates after treatment with systemic neutralizing antibodies. Cell Host Microbe 29, 551–563 e555, doi:10.1016/j.chom.2021.02.019 (2021).

2 Zhu, N. et al. A Novel Coronavirus from Patients with Pneumonia in China, 2019. N Engl J Med 382, 727–733, doi:10.1056/NEJMoa2001017 (2020).

3 Johns Hopkins University. The Center for Systems Science and Engineering (CSSE) COVID-19 Dashboard, https://gisanddata.maps.arcgis.com/apps/dashboards/bda7594740fd40299423467b48e9ecf6 (2021).

4 Weinreich, D. M. et al. REGN-COV2, a Neutralizing Antibody Cocktail, in Outpatients with Covid-19. N Engl J Med 384, 238–251, doi:10.1056/NEJMoa2035002 (2021).

5 Chen, P. et al. SARS-CoV-2 Neutralizing Antibody LY-CoV555 in Outpatients with Covid-19. N Engl J Med 384, 229–237, doi:10.1056/NEJMoa2029849 (2021).

6 Hamers-Casterman, C. et al. Naturally occurring antibodies devoid of light chains. Nature 363, 446–448, doi:10.1038/363446a0 (1993).

7 Muyldermans, S. Nanobodies: natural single-domain antibodies. Annu Rev Biochem 82, 775–797, doi:10.1146/annurev-biochem-063011-092449 (2013).

8 Pardon, E. et al. A general protocol for the generation of Nanobodies for structural biology. Nat Protoc 9, 674–693, doi:10.1038/nprot.2014.039 (2014).

9 Esparza, T. J., Martin, N. P., Anderson, G. P., Goldman, E. R. & Brody, D. L. High affinity nanobodies block SARS-CoV-2 spike receptor binding domain interaction with human angiotensin converting enzyme. Sci Rep 10, 22370, doi:10.1038/s41598-020-79036-0 (2020).

10 Koenig, P. A. et al. Structure-guided multivalent nanobodies block SARS-CoV-2 infection and suppress mutational escape. Science 371, doi:10.1126/science.abe6230 (2021).

11 Xu, J. et al. Nanobodies from camelid mice and llamas neutralize SARS-CoV-2 variants. Nature 595, 278–282, doi:10.1038/s41586-021-03676-z (2021).

12 Xiang, Y. et al. Versatile and multivalent nanobodies efficiently neutralize SARS-CoV-2. Science 370, 1479–1484, doi:10.1126/science.abe4747 (2020).

13 Pymm, P. et al. Nanobody cocktails potently neutralize SARS-CoV-2 D614G N501Y variant and protect mice. Proc Natl Acad Sci U S A 118, doi:10.1073/pnas.2101918118 (2021).

14 Li, T. et al. A synthetic nanobody targeting RBD protects hamsters from SARS-CoV-2 infection. Nat Commun 12, 4635, doi:10.1038/s41467-021-24905-z (2021).

15 Guttler, T. et al. Neutralization of SARS-CoV-2 by highly potent, hyperthermostable, and mutation-tolerant nanobodies. EMBO J, e107985, doi:10.15252/embj.2021107985 (2021).

16 Schoof, M. et al. An ultrapotent synthetic nanobody neutralizes SARS-CoV-2 by stabilizing inactive Spike. Science 370, 1473–1479, doi:10.1126/science.abe3255 (2020).

17 Hoffmann, M. et al. SARS-CoV-2 Cell Entry Depends on ACE2 and TMPRSS2 and Is Blocked by a Clinically Proven Protease Inhibitor. Cell 181, 271–280 e278, doi:10.1016/j.cell.2020.02.052 (2020).

18 Letko, M., Marzi, A. & Munster, V. Functional assessment of cell entry and receptor usage for SARS-CoV-2 and other lineage B betacoronaviruses. Nat Microbiol 5, 562–569, doi:10.1038/s41564-020-0688-y (2020).

19 Martin, A. C. Accessing the Kabat antibody sequence database by computer. Proteins 25, 130–133, doi:10.1002/(SICI)1097-0134(199605)25:1<130::AID-PROT11>3.0.CO;2-L (1996).

20 Rappazzo, C. G. et al. Broad and potent activity against SARS-like viruses by an engineered human monoclonal antibody. Science 371, 823–829, doi:10.1126/science.abf4830 (2021).

21 Yao, H. et al. A high-affinity RBD-targeting nanobody improves fusion partner’s potency against SARS-CoV-2. PLoS Pathog 17, e1009328, doi:10.1371/journal.ppat.1009328 (2021).

22 Barnes, C. O. et al. SARS-CoV-2 neutralizing antibody structures inform therapeutic strategies. Nature 588, 682–687, doi:10.1038/s41586-020-2852-1 (2020).

23 Webb, N. E., Montefiori, D. C. & Lee, B. Dose-response curve slope helps predict therapeutic potency and breadth of HIV broadly neutralizing antibodies. Nat Commun 6, 8443, doi:10.1038/ncomms9443 (2015).

24 Van Heeke, G. et al. Nanobodies(R) as inhaled biotherapeutics for lung diseases. Pharmacol Ther 169, 47–56, doi:10.1016/j.pharmthera.2016.06.012 (2017).

25 Larios Mora, A. et al. Delivery of ALX-0171 by inhalation greatly reduces respiratory syncytial virus disease in newborn lambs. MAbs 10, 778–795, doi:10.1080/19420862.2018.1470727 (2018).

26 Detalle, L. et al. Generation and Characterization of ALX-0171, a Potent Novel Therapeutic Nanobody for the Treatment of Respiratory Syncytial Virus Infection. Antimicrob Agents Chemother 60, 6–13, doi:10.1128/AAC.01802-15 (2016).

27 Imai, M. et al. Syrian hamsters as a small animal model for SARS-CoV-2 infection and countermeasure development. Proc Natl Acad Sci U S A 117, 16587–16595, doi:10.1073/pnas.2009799117 (2020).

28 Huo, J. et al. A potent SARS-CoV-2 neutralising nanobody shows therapeutic efficacy in the Syrian golden hamster model of COVID-19. Nat Commun 12, 5469, doi:10.1038/s41467-021-25480-z (2021).

29 Li, W. et al. High Potency of a Bivalent Human VH Domain in SARS-CoV-2 Animal Models. Cell 183, 429–441 e416, doi:10.1016/j.cell.2020.09.007 (2020).

30 Ye, G. et al. The Development of Nanosota-1 as anti-SARS-CoV-2 nanobody drug candidates. Elife 10, doi:10.7554/eLife.64815 (2021).

31 Nambulli, S. et al. Inhalable Nanobody (PiN-21) prevents and treats SARS-CoV-2 infections in Syrian hamsters at ultra-low doses. Sci Adv 7, doi:10.1126/sciadv.abh0319 (2021).

32 Moliner-Morro, A. et al. Picomolar SARS-CoV-2 Neutralization Using Multi-Arm PEG Nanobody Constructs. Biomolecules 10, doi:10.3390/biom10121661 (2020).

33 Miller, A., Carr, S., Rabbitts, T. & Ali, H. Multimeric antibodies with increased valency surpassing functional affinity and potency thresholds using novel formats. MAbs 12, 1752529, doi:10.1080/19420862.2020.1752529 (2020).

34 Zupancic, J. M. et al. Directed evolution of potent neutralizing nanobodies against SARS-CoV-2 using CDR-swapping mutagenesis. Cell Chem Biol 28, 1379–1388 e1377, doi:10.1016/j.chembiol.2021.05.019 (2021).

35 Huo, J. et al. Neutralizing nanobodies bind SARS-CoV-2 spike RBD and block interaction with ACE2. Nat Struct Mol Biol 27, 846–854, doi:10.1038/s41594-020-0469-6 (2020).

36 Minor, W., Cymborowski, M., Otwinowski, Z. & Chruszcz, M. HKL-3000: the integration of data reduction and structure solution--from diffraction images to an initial model in minutes. Acta Crystallogr D Biol Crystallogr 62, 859–866, doi:10.1107/S0907444906019949 (2006).

37 Winn, M. D. et al. Overview of the CCP4 suite and current developments. Acta Crystallogr D Biol Crystallogr 67, 235–242, doi:10.1107/S0907444910045749 (2011).

38 Emsley, P. & Cowtan, K. Coot: model-building tools for molecular graphics. Acta Crystallogr D Biol Crystallogr 60, 2126–2132, doi:10.1107/S0907444904019158 (2004).

39 Liebschner, D. et al. Macromolecular structure determination using X-rays, neutrons and electrons: recent developments in Phenix. Acta Crystallogr D Struct Biol 75, 861–877, doi:10.1107/S2059798319011471 (2019).

40 Chen, V. B. et al. MolProbity: all-atom structure validation for macromolecular crystallography. Acta Crystallogr D Biol Crystallogr 66, 12–21, doi:10.1107/S0907444909042073 (2010).

41 Krissinel, E. & Henrick, K. Inference of macromolecular assemblies from crystalline state. J Mol Biol 372, 774–797, doi:10.1016/j.jmb.2007.05.022 (2007).

42 Wilkins, M. R. et al. Protein identification and analysis tools in the ExPASy server. Methods Mol Biol 112, 531–552, doi:10.1385/1-59259-584-7:531 (1999).

43 Chen, S. H. et al. Production of Viral Constructs for Neuroanatomy, Calcium Imaging, and Optogenetics. Curr Protoc Neurosci 87, e66, doi:10.1002/cpns.66 (2019).

44 Bosco-Lauth, A. M. et al. Experimental infection of domestic dogs and cats with SARS-CoV-2: Pathogenesis, transmission, and response to reexposure in cats. Proc Natl Acad Sci U S A 117, 26382–26388, doi:10.1073/pnas.2013102117 (2020).

45 Jia, Q. et al. Replicating bacterium-vectored vaccine expressing SARS-CoV-2 Membrane and Nucleocapsid proteins protects against severe COVID-19-like disease in hamsters. NPJ Vaccines 6, 47, doi:10.1038/s41541-021-00321-8 (2021).

46 Walls, A. C. et al. Structure, Function, and Antigenicity of the SARS-CoV-2 Spike Glycoprotein. Cell 183, 1735, doi:10.1016/j.cell.2020.11.032 (2020).

47 Wrapp, D. et al. Cryo-EM structure of the 2019-nCoV spike in the prefusion conformation. Science 367, 1260–1263, doi:10.1126/science.abb2507 (2020).

