## Supplemental Material for "Nebulized delivery of a broadly neutralizing SARS-CoV-2 RBD-specific nanobody prevents clinical, virological and pathological disease in a Syrian hamster model of COVID-19"

### Supplementary Results

#### *In vitro* affinity measurements of NIH-CoVnb-112 binding to SARS-CoV-2 RBD variants

Additional bio-layer interferometry was performed to further evaluate the potential impact of single RBD mutations within and adjacent to the binding epitope of NIH-CoVnb-112. The assays were conducted using the same parameters as the previous measurements. Using SARS-CoV-2 RBDs with single residue point mutations at K417N, L452R, T478K, and E484K, biosensor bound NIH-CoVnb-112 was used to measure the $k_{on}$ and $k_{off}$ values as described. Curve fitting produced $K_D$ values for E484K: 1.38nM (**Supplementary Fig. 1a**), K417N: 1.42nM (**Supplementary Fig. 1b**), L452R: 4.88nM (**Supplementary Fig. 1c**), and T478K: 0.497nM (**Supplementary Fig. 1d**). The individual $k_{on}$ values (**Supplementary Table 2**) range from $1.20 \times 10^5$ to $4.10 \times 10^5$ $M^{-1}s^{-1}$ or ~0.5 to 2-fold of the prototype RBD, indicating robust association independent of residue mutation. Similarly, three of the four variant RBDs had dissociation constants ($k_{off}$) which were comparably ranging from $1.27 \times 10^{-4}$ to $3.04 \times 10^{-4}$ $s^{-1}$. However, a 5-fold increase in dissociation was observed for the L452R mutation with a $k_{off}$ of $1.13 \times 10^{-3}$ $s^{-1}$. This data is consistent with our structural prediction that L452R mutation in Delta strain may cause steric hindrance with the CDR3 of NIH-CoVnb-112, although another mutation T478K appears to introduce a salt-bridge with the $D^{61}$ on CDR2. (**Fig. 2b**). Interestingly, the counterbalance between the weaker binding to the L452R and stronger binding to the T478K mutations ultimately results in a similar equilibrium dissociation constant for the Delta strain RBD relative to the prototype sequence.

#### Pilot study of nebulization treatment of NIH-CoVnb-112 in a hamster model of COVID-19

Previously[1], we utilized an Aerogen Solo mesh nebulizer for in-vitro characterization of NIH-CoVnb-112. For the study described here, the PARI Sprint nebulizer was an existing component of the aerosol delivery system. Thus, to ensure stability post-nebulization, we used the PARI Sprint nebulizer in-line with our custom collection system to nebulize a 2mg/mL solution of NIH-CoVnb-112. Following recovery, the concentration of the nebulized nanobody was measured by absorbance and determined a 96% recovery of input protein. Pre-nebulization and post-nebulization samples were injected onto a Superdex 75 Increase 10/300 GL SEC column and the resulting absorbance profiles were compared. We observe near perfect overlap of the two samples (**Supplementary Fig. 2**), with no apparent aggregation or degradation of protein, indicating excellent stability using the PARI Sprint system.

Confident in the post-nebulization nanobody stability, we then explored the potential for *in vivo* delivery and therapeutic potential of NIH-CoVnb-112 in a Syrian hamster model of COVID-19. Adult male Syrian hamsters, 12 weeks of age, were acclimated to the ABSL3 facility prior to use and monitored for overt health concerns. For the initial study, we chose to assess nanobody prophylaxis (-24hr. pre infection) and post-exposure (+8hr. post-infection) treatment as outlined (**Supplementary Fig. 3a**). Cohorts of 6 hamsters were randomized into the following groups: (1) saline/saline, (2) saline/nanobody, (3) nanobody/saline, (4) nanobody/nanobody, (5) no treatment control. Twenty-four hours before the virus challenge, groups of 3 animals were placed in a containment box with attached nebulizer, animals received nebulized exposure of a 5mL solution containing normal saline alone or normal saline with 10mg/mL NIH-CoVnb-112 for 20mins. Twenty-four hours after the initial dose, under light anesthesia, all animals were challenged with ~$10^4$ pfu of SARS-CoV-2 (2019-nCoV/USA-WA1/2020 strain) by intranasal administration. A second nebulization was given at 8hrs. following virus challenge as indicated (**Supplementary Fig. 3a**). Animals were observed for weight on days 0 - 7 post infection (**Supplementary Fig. 3b**) and oropharyngeal swabs taken on days 1, 2, and 3 post infection (**Supplementary Fig. 4**). Half of each group was euthanized on day 3 post infection and the remaining half euthanized on day 7 post infection for viral titer and histopathology assessments.

It was previously reported that Syrian hamsters infected with SARS-CoV-2 undergo transient weight loss and severe lung pathology during the subacute phase of infection.[25] Body weight measurements for all groups (**Supplementary Fig. 3b**) exhibited a negative trend throughout the time course of the study. In both nasal turbinates and cranial lung tissue harvested at 3 days post infection (**Supplementary Fig. 3c-d**), there was a sustained level of viral burden, as measured by double-overlay plaque assay, with no apparent difference between treatment groups. In the oropharyngeal swabs (**Supplementary Fig. 4**), all groups followed a day-associated decrease in viral titer with no apparent difference between groups. Lung and tracheal tissue were collected at day 3 and 7 post infection to assess the impact of nebulization delivery of NIH-CoVnb-112 on pathology. A veterinary pathologist, blinded to status, performed histopathological scoring on H&E sections using an established semi-quantitative lung scoring scale (**Supplementary Table 4, Supplementary Fig. 3e-f, and Supplementary Fig. 5**).[2] Histological observation of the saline/saline and no treatment controls revealed typical histological signs of pulmonary disease, including perivascular and peribronchial mononuclear cell infiltration, focal tracheal necrosis and exfoliation, and moderate to severe septal and intra-alveolar infiltration of mononuclear cells. In the saline/nanobody and nanobody/saline groups the presence of peribronchial mononuclear cell infiltration was milder, the tracheal epithelium was intact, and perivascular infiltration reduced in the saline/nanobody group compared to the nanobody/saline group. Both groups exhibited a reduction in day 3 intra-alveolar infiltration yet progressed to marked interstitial and intra-alveolar infiltration of mononuclear cells by day 7. By contrast, the nanobody/nanobody treatment group was largely within normal limits at both time points, with mild tracheal leukocyte infiltration on day 3, mild bronchial epithelial hypertrophy on day 7, and mild alveolar mononuclear cell infiltration on day 7. The total lung score and subset scores (**Supplementary Fig. 3e-f and Supplementary Fig. 6**) reveal an early reduction in lung score for the nanobody/nanobody (day 3 mean score = 8.3) versus the saline/saline and no treatment groups (day 3 mean score = 26.3 and 30, respectively). By day 7 there was a global increase in all groups, with the nanobody/nanobody group (day 7 mean score = 23.7) maintaining an overall lower score than the control groups (day 7 mean score = 41.7 and 40.7, respectively). The saline/nanobody and nanobody/saline groups had lung scores (day 3 mean score = 19.3 and 22, day 7 mean score = 34.3 and 36, respectively) modestly lower than the control groups. Thus, the pilot study of nebulization delivery of NIH-CoVnb-112 demonstrated feasibility and some indications of potential therapeutic efficacy.

**Supplementary Table 1. Crystallographic data collection and refinement statistics.**

| | NIH-CoVnb-112 and SARS-CoV-2 |
|---|---|
| **Data collection** | |
| Wavelength, Å | 0.979 |
| Resolution range, Å | 54.2 - 2.82 (2.921 - 2.82) |
| Space group | $P2_1$ |
| Unit cell parameter | |
| a, b, c, Å | 32.7, 59.0, 216.9 |
| α, β, γ, ° | 90.0, 91.3, 90.0 |
| Redundancy | 8.08 (8.16) |
| Completeness, % | 79.72 (74.94) |
| Mean I/sigma(I) | 22.59 (4.98) |
| $R_{merge}$[a] | 0.150 (0.314) |
| $R_{pim}$[b] | 0.100 (0.213) |
| $CC_{1/2}$[c] | 0.944 (0.791) |
| Wilson $B_{factor}$, ($1/Å^2$)[d] | 22.80 |
| **Refinement** | |
| $R_{work}$[e] | 0.247 (0.334) |
| $R_{free}$[f] | 0.2957 (0.4123) |
| Resolution, Å | 54.2 – 2.82 |
| # of non-hydrogen | |
| proteins | 5062 |
| Water | 8 |
| Overall $B_{factor}$, ($Å^2$) | |
| Proteins | 16.78 |
| Ligands | 74.43 |
| Water | 16.55 |
| RMS (bond lengths), Å | 0.012 |
| RMS (bond angles), ° | 1.44 |
| Ramachandran[g] | |
| Favored, % | 93.93 |
| Allowed, % | 6.07 |
| Outliers, % | 0.0 |
| PDB ID | 7RBY |

Statistics for the highest-resolution shell are shown in parentheses.

[a]$R_{merge} = \sum | I - \langle I \rangle | / \sum I$, where $I$ is the observed intensity and $\langle I \rangle$ is the average intensity obtained from multiple observations of symmetry-related reflections after rejections

[b]$R_{pim}$ = as defined in Weiss.[3]

[c]$CC_{1/2}$ = as defined by Karplus and Diederichs.[4]

[d]Wilson $B_{factor}$ as calculated in Popov and Bourenkov.[5]

[e]$R = \sum \| F_o | - | F_c \| / \sum | F_o |$, where $F_o$ and $F_c$ are the observed and calculated structure factors, respectively

[f]$R_{free}$ = as defined by Brünger et al.[6]

[g]Calculated with MolProbity.[7]

### Supplementary Table 2. NIH-CoVnb-112 Affinity Binding Values Against SARS-CoV-2 Variant RBDs

| SARS-CoV-2 RBD | $k_{on}$(1/Ms) | $k_{dis}$(1/s) | KD (M) |
|---|---|---|---|
| **Prototype** | $2.23x10^{5}$ | $2.36x10^{-4}$ | $1.59x10^{-9}$ |
| **Alpha (B.1.1.7) N501Y** | $1.53x10^{5}$ | $4.28x10^{-4}$ | $3.00x10^{-9}$ |
| **Beta (B.1.351)** | $2.01x10^{5}$ | $7.05x10^{-4}$ | $4.28x10^{-9}$ |
| **Gamma (P.1)** | $2.14x10^{5}$ | $8.22x10^{-4}$ | $4.16x10^{-9}$ |
| **Delta (B.1.617.2)** | $4.61x10^{5}$ | $6.09x10^{-4}$ | $1.66x10^{-9}$ |
| **K417N** | $1.20x10^{5}$ | $1.57x10^{-4}$ | $1.42x10^{-9}$ |
| **E484K** | $2.56x10^{5}$ | $3.04x10^{-4}$ | $1.38x10^{-9}$ |
| **L452R** | $2.39x10^{5}$ | $1.13x10^{-3}$ | $4.88x10^{-9}$ |
| **T478K** | $4.10x10^{5}$ | $1.27x10^{-4}$ | $4.97x10^{-10}$ |

### Supplementary Table 3. Pseudovirus neutralization assay curve fit parameters

| Parameter | Prototype | Alpha - B.1.1.7 | Beta - B.1.351 | Gamma - P.1 | Delta - B.1.612.7 |
|---|---|---|---|---|---|
| **Top (%)** | 99.57 | 92.39 | 100.4 | 100.1 | 86.06 |
| **Bottom (%)** | 3.912 | -3.066 | -8.574 | 6.215 | 3.256 |
| **$EC_{50}$ (nM)** | 14.27 | 9.364 | 15.82 | 17.61 | 14.52 |
| **HillSlope** | 1.155 | 0.9714 | 1.199 | 1.246 | 1.162 |

**Supplementary Table 4. Histopathological scoring of two dose nebulization delivered NIH-CoVnb-112 and saline combinations**

| Treatment group | Trachea | Bronchiole | Blood vessel | Alveolar parenchyma |
|---|---|---|---|---|
| **Saline/Saline 3dpi** | Focal epithelial cell degeneration, necrosis, and exfoliation (arrow). Mild infiltration of granulocytes (heterophils; the hamster equivalent of neutrophils) and mononuclear leukocytes. | Moderate to severe infiltration of monocytes & heterophils with epithelial transmigration and exfoliation of epithelial cells. | Severe mural infiltration of monocytes and heterophils disrupting the vessel intima and media. | Focal, moderate alveolar septal (interstitial) & intra-alveolar infiltration of mononuclear cells (MNC) and heterophils (area indicated with star). |
| **Saline/Saline 7dpi** | Histologically within normal limits (WNL) | Moderate epithelial hypertrophy & minimal leukocyte infiltration | Mild endothelial hypertrophy, but vessels otherwise WNL | Moderate to severe interstitial & intra-alveolar leukocyte infiltration. Mild erythrocyte diapedesis. Severe hypertrophy of of pneumocytes. |
| **Saline/Nanobody 3dpi** | Mild mucosal infiltration of MNC. Epithelium intact. | Mild to moderate infiltration of MNC & heterophils, mild epithelial cells degeneration and detachment (exfoliation). | Mild leukocyte margination (arrow) and transmigration. Mild endothelial cell hypertrophy. | Mild to moderate interstitial & intra-alveolar leukocyte infiltration. |
| **Saline/Nanobody 7dpi** | Mild mucosal infiltration of MNC. Epithelium intact. | Moderate epithelial hypertrophy & minimal leukocyte infiltration | Mild perivascular infiltration of MNC (arrow). Vessels otherwise WNL | Marked interstitial & intra-alveolar leukocyte infiltration & pneumocyte hypertrophy. |
| **Nanobody/Saline 3dpi** | Severe epithelial cell degeneration, necrosis & exfoliation (arrow) accompanied by MNC & heterophil infiltration. | Moderate to severe infiltration of monocytes & heterophils with epithelial transmigration and exfoliation of epithelial cells. | Moderate mural infiltration of MNC & heterophils variably disrupting intima & media (arrows). | Mild alveolar septal infiltration (stars) of mainly MNC. |
| **Nanobody/Saline 7dpi** | Minimal leukocyte infiltration in subepithelial interstitium. Epithelium intact | Moderate epithelial hypertrophy & minimal leukocyte infiltration. | Moderate to severe transmural infiltration of MNC (arrows) with endothelial hypertrophy and intimal hyperplasia (repair/scarring). | Marked interstitial & intra-alveolar infiltration of MNC and heterophils, and pneumocyte hypertrophy. |
| **Nanobody/Nanobody 3dpi** | Mild leukocyte infiltration in subepithelial interstitium and intraepithelial migration, but epithelium is intact. | WNL | WNL | WNL |
| **Nanobody/Nanobody 7dpi** | WNL | Mild epithelial hypertrophy. | WNL | Mild thickening of alveolar septae (star) due to mild MNC infiltration. |
| **No nebulization ctrl. 3dpi** | Severe degeneration & necrosis with exfoliation of epithelial cells (arrow) accompanied by infiltration of MNC and heterophils. | Severe infiltration and epithelial transmigration of heterophils and monocytes, exfoliation of degenerating epithelial cells and accumulation of cellular debris in lumen (star). | Moderate transmural migration (arrow) and infiltration of monocytes & heterophils | Moderate to severe septal and intra-alveolar infiltration of monocytes and heterophils accompanied by erythrocyte diapedesis or frank hemorrhage. |
| **No nebulization ctrl. 7dpi** | Mild mucosal infiltration of MNC. Epithelium intact. | Moderate epithelial hypertrophy & minimal leukocyte infiltration | Moderate transmural migration (arrow) and infiltration of monocytes & heterophils | Marked interstitial & intra-alveolar infiltration of MNC and heterophils, and pneumocyte hypertrophy. |

**WNL**: within normal limits; **MNC**: mononuclear cell; **dpi**: days post-infection

### Supplementary Table 5. Histopathological scoring of multi dose nebulization delivered NIH-CoVnb-112 versus saline control

| Treatment group | Trachea | Bronchiole | Blood vessel | Alveolar parenchyma |
|---|---|---|---|---|
| **Nanobody (3dose) 3dpi** | WNL | WNL | WNL | WNL |
| **Nanobody (4dose) 7dpi** | Minimal MNC infiltration in subepithelial interstitium (arrow). Epithelium intact. | WNL | Minimal perivascular accumulation of MNC (arrow). | WNL |
| **Saline (3dose) 3dpi** | Minimal MNC infiltration in subepithelial interstitium and intraepithelial migration, but epithelium intact. | Moderate to severe peribronchial and transepithelial leukocyte infiltration, epithelial cell exfoliation and cellular debris accumulating in the lumen (arrow). | Very severe transmural MNC & heterophil infiltration with very severe accumulation of leukocytes in intima and between smooth muscle cell layers in the media (arrow). Severe perivascular edema (star). | Severe inter- & intra-alveolar MNC and heterophil infiltration, erythrocyte diapedesis and frank hemorrhage occluding alveolar spaces. |
| **Saline (4dose) 7dpi** | Mild MNC infiltration in subepithelial interstitium. Epithelium intact | Pronounced contraction of bronchiole, epithelial hypertrophy and moderate leukocyte infiltration in peribronchial interstitium with transepithelial migration | Moderate transmural leukocyte migration and perivascular accumulation (arrow). | Very severe intra- and interalveolar leukocyte infiltration and pneumocyte hypertrophy causing complete effacement of normal alveolar morphology. |

**WNL**: within normal limits; **MNC**: mononuclear cell; **dpi**: days post-infection

**Supplementary Fig. 1**

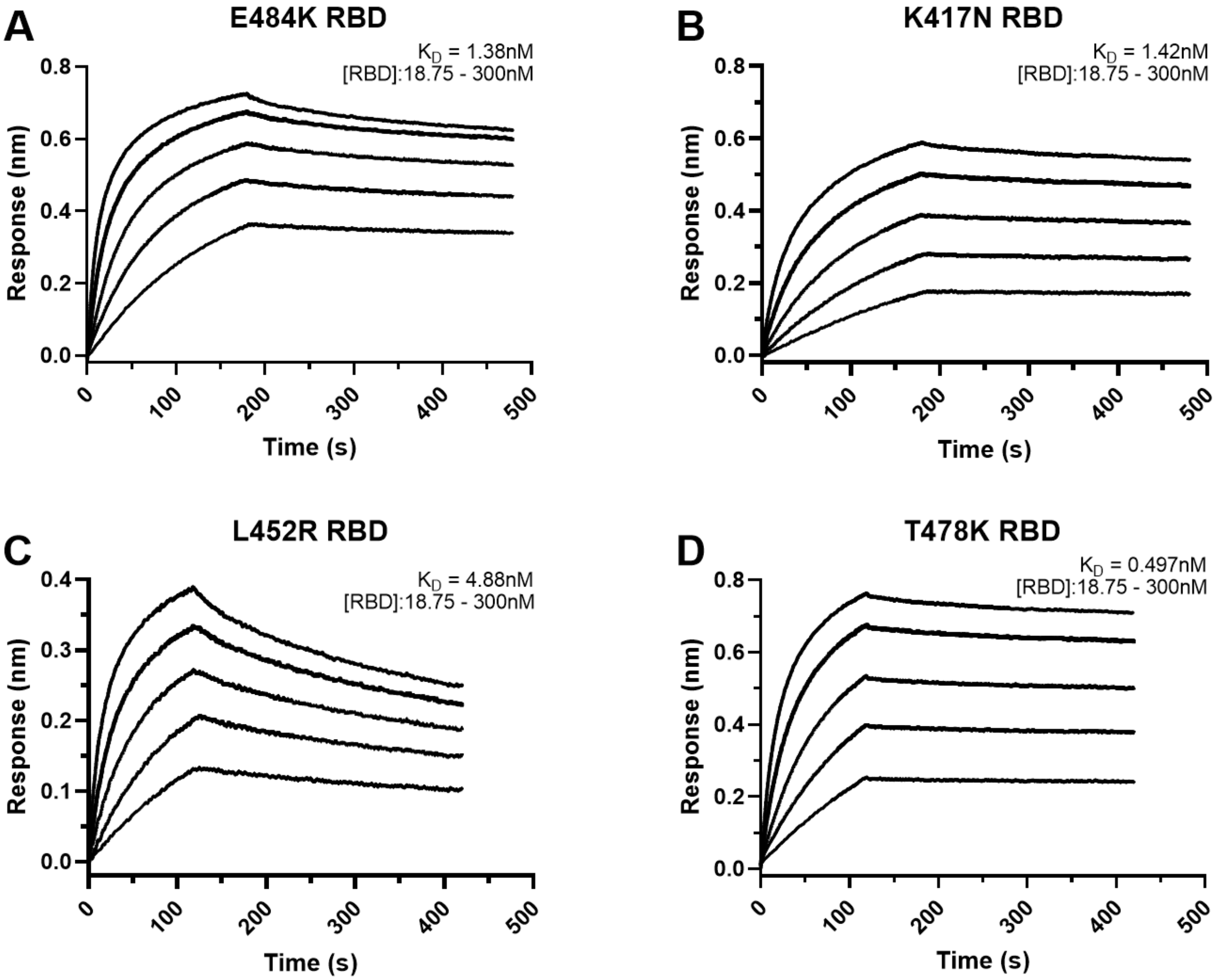

**Supplementary Fig. 1. Affinity binding curves of NIH-CoVnb-112 against SARS-CoV-2 RBD variant single point mutations**. Using Biolayer Interferometry on a BioForte Octet Red96 system, association and dissociation rates were determined by immobilizing biotinylated-NIH-CoVnb-112 onto streptavidin coated optical sensors (**a-d**). The nanobody-bound sensors were incubated with a concentration range (18.75 – 300nM) of recombinant single point mutation RBDs, (**a**) E484K, (**b**) K417N, (**c**) L452R, and (**d**) T478K, for a set time interval to allow association. The sensors were then moved to RBD-free solution and allowed to dissociate over a time interval. Curve fitting using a 1:1 interaction model allows for the affinity constant ($K_D$) to be measured for each RBD variant as detailed in **Supplementary Table 2**.

**Supplementary Fig. 2**

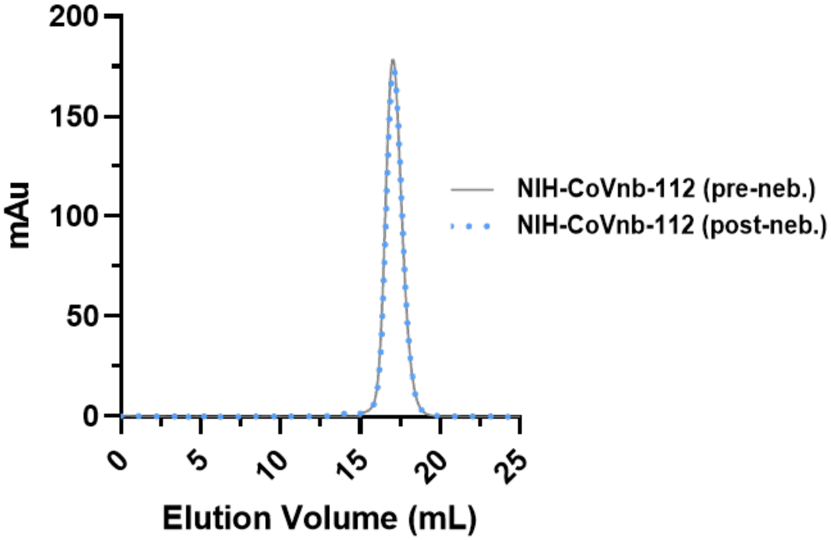

**Supplementary Fig. 2. Pre- and post-nebulization effect on NIH-CoVnb-112 using a PARI Sprint nebulizer.** A PARI Sprint nebulizer was used to aerosolize a 2mg/mL solution of NIH-CoVnb-112 and collected using a custom condenser. Following nebulization and recovery, an equal volume of the pre- and post-nebulization material was injected on a Superdex 75 10/300 GL on an AKTA Purifier. Size exclusion chromatography reveals a prominent overlapping peak in both pre- and post-nebulization samples. The overlapping and symmetrical peaks indicate uniformity and lack of aggregation or degradation of the nanobody during nebulization.

**Supplementary Fig. 3**

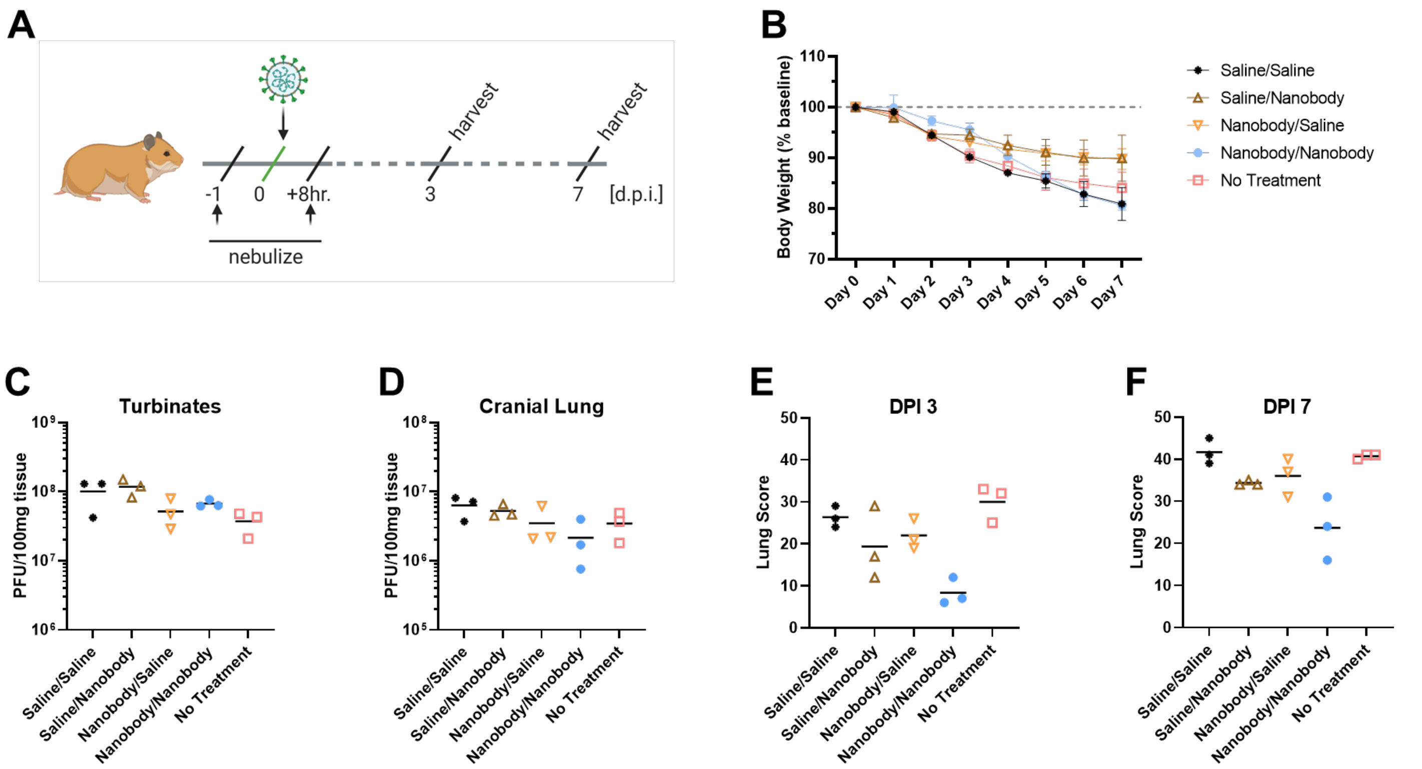

**Supplementary Fig. 3. NIH-CoVnb-112 nebulization treatment in a hamster model of SARS-CoV-2**. (**a**) Schematic overview of study design for nebulization treatment of Syrian hamsters with NIH-CoVnb-112. Adult Syrian hamsters (n=3/group, males, 12 weeks old) were treated at -24hr prior to virus challenge with nebulized saline or NIH-CoVnb-112 (10mg/mL) in a 5mL volume over 20mins. Following intranasal challenge, each group was treated at 8hrs post infection with the same condition. One group received no treatment. Animals were weighed daily, and oral swabs taken at dpi 1-3. Groups were euthanized at day 3 and day 7 post infection respectively and sample taken for assessment. (Figure elements generated using BioRender.com) (**b**) Body weight change as a percentage of baseline weight for day 7 post infection group (n=3/group, mean +/- SD) (**c**) Viral burden from turbinate tissue on day 3 post infection groups. (d) Viral burden from cranial lung tissue of day 3 post infection groups. (**e-f**) Summed pulmonary lesion scores from hematoxylin and eosin-stained tissue sections collected on day 3 and day 7 post-infection. Metrics included overall lesion extent, bronchitis, alveolitis, pneumocyte hyperplasia, vasculitis, and interstitial inflammation; each on a 0–4 or 0–5 scale and scores summed for day 3 and day 7 post infection are described in **Supplementary Fig. 6**.

**Supplementary Fig. 4**

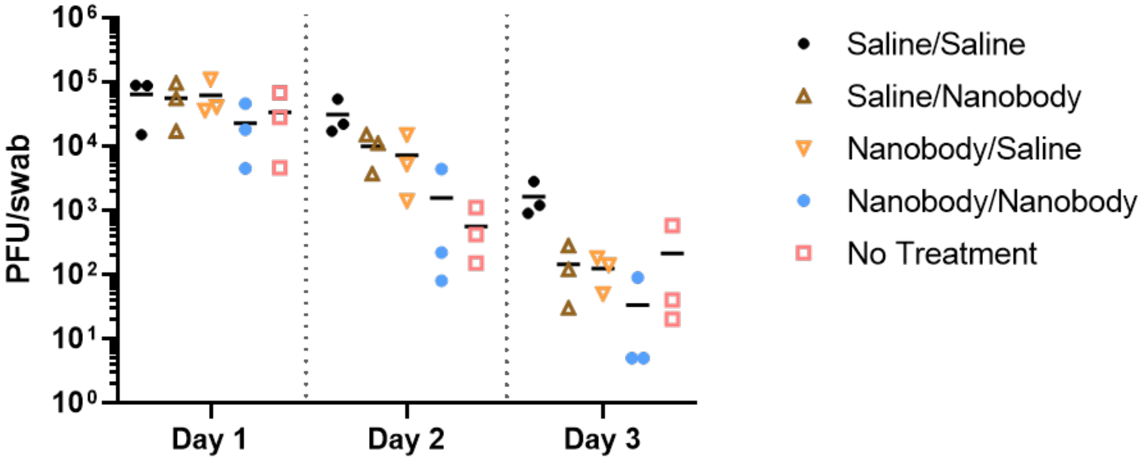

**Supplementary Fig. 4. Oropharyngeal swab SARS-CoV-2 viral burden from two dose nebulization treatment assessment.** Oral swabs taken on day 1, 2, and 3 post infection were quantified by double-overlay plaque assay on Vero E6 cells. At 48–72hr post-infection, plaques were viewed with a lightbox counted as plaque forming units (PFU) per swab sample.

**Supplementary Fig. 5**

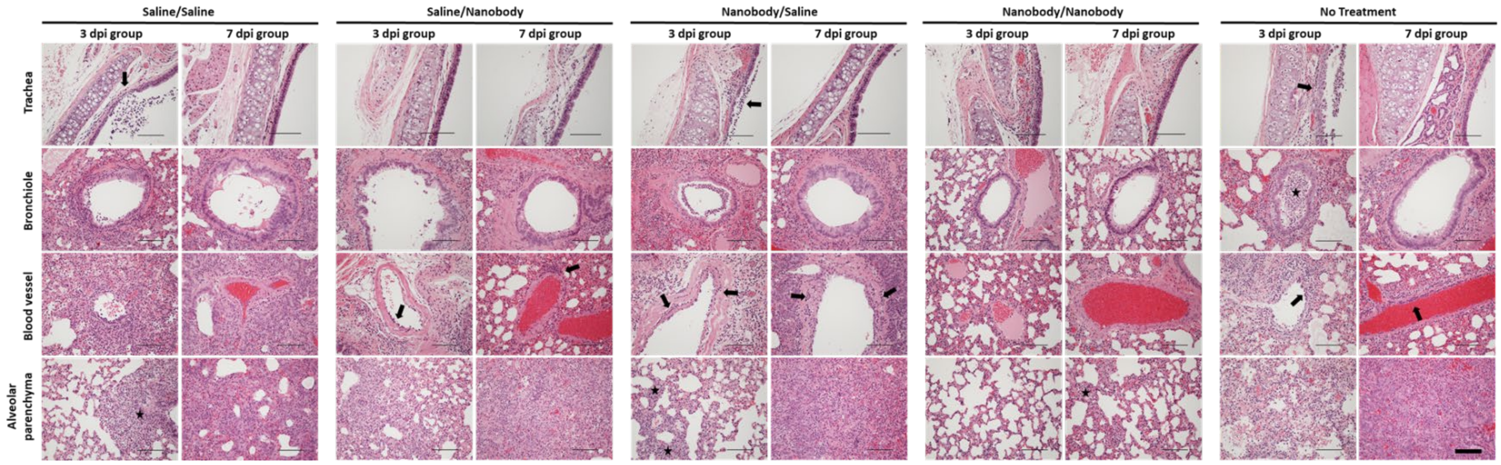

**Supplementary Fig. 5. Histopathology of trachea and lung from two dose nebulization treated Syrian hamsters following SARS-CoV-2 infection**. Representative hematoxylin and eosin-stained tissue sections from day 3 and day 7 post infection groups (n = 4 per group) for trachea, bronchioles, blood vessels, and alveolar parenchyma. Corresponding pathologist indications and notations found in **Supplementary Table 4**; arrows and stars represent pathological hallmarks as described in the table. Images acquired at 200x magnification. Scale bars for all images = 300µm.

**Supplementary Fig. 6**

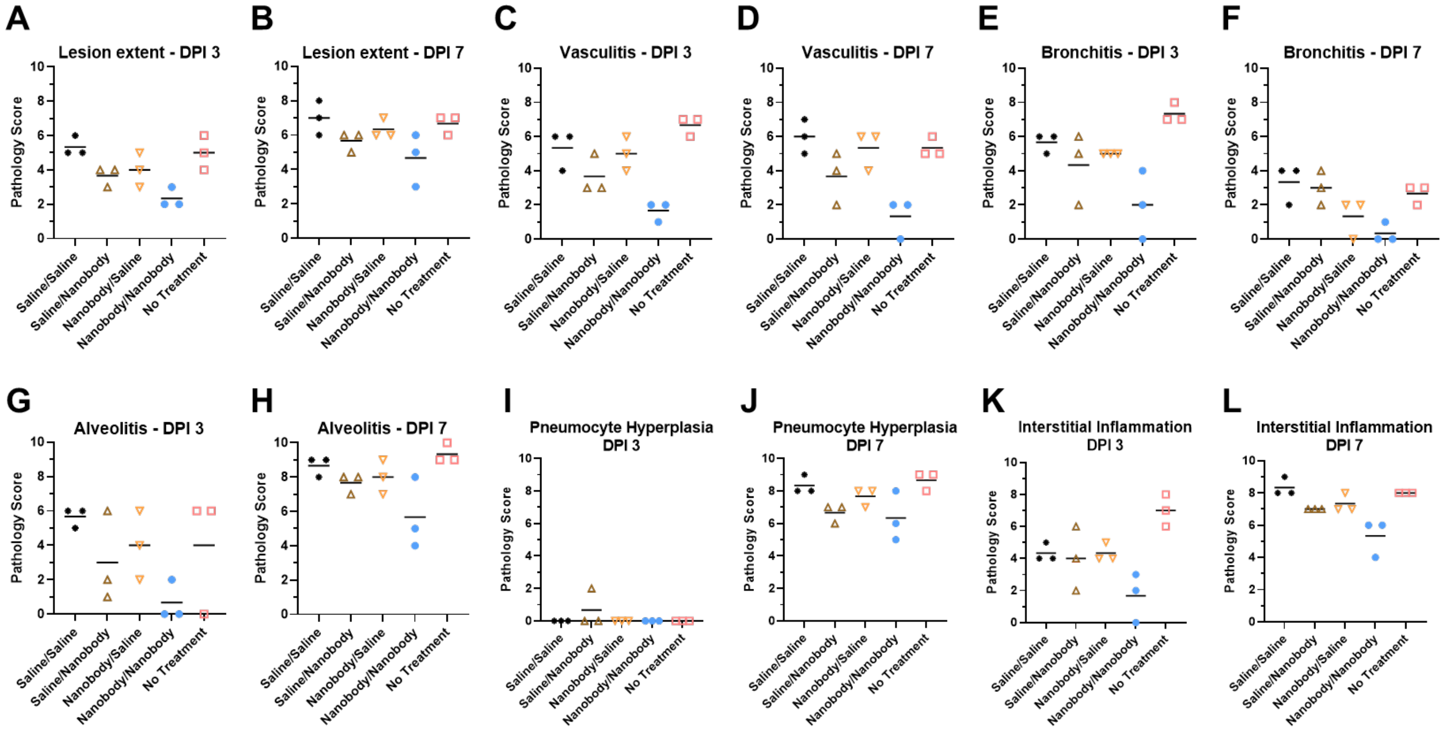

**Supplementary Fig. 6. Histological lung score individual metrics for two dose nebulization treated hamsters.** Hematoxylin and eosin-stained tissue sections from the day 3 and day 7 post infection animals were semi-quantitively scored by a pathologist blinded to condition using an established scoring metric. Metrics included overall (**a,b**) lesion extent, (**c,d**) vasculitis, (**e,f**) bronchiolitis, (**g,h**) alveolitis, (**i,j**) pneumocyte hyperplasia, and (**k,l**) interstitial inflammation; each on a 0–4 or 0–5 scale.

**Supplementary Fig. 7**

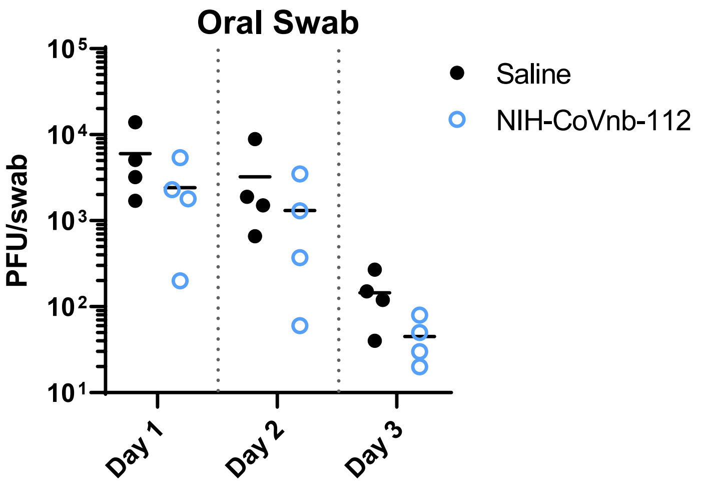

**Supplementary Fig. 7. Oropharyngeal swab SARS-CoV-2 viral burden from multi dose nebulization treatment assessment.** Oral swabs taken on day 1, 2, and 3 post infection were quantified by double-overlay plaque assay using Vero E6 cells. Swabs were processed in Tris-buffered Minimum Essential Medium (MEM) with 1% BSA and inoculated onto confluent monolayers of Vero E6 cells seeded in 6-well cell culture plates. At 48–72hr post-infection, plaques were viewed with a lightbox counted as plaque forming units (PFU) per swab sample.

**Supplementary Fig. 8**

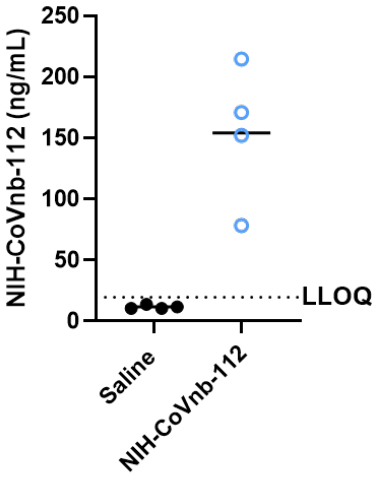

**Supplementary Fig. 8. Quantification of NIH-CoVnb-112 in bronchoalveolar lavage fluid following nebulization delivery in hamster.** BALF collected on day 3 post infection and 24hrs following the third nebulization dose of nanobody. Using an ELISA format which immobilized SARS-CoV-2 RBD onto the plate surface, nanobody was detected following binding to RBD using an anti-alpaca peroxidase conjugated secondary monoclonal. A standard curve using NIH-CoVnb-112 allowed for interpolation of the BALF samples. The lower limit of quantification (LLOQ) of the assay (19.5ng/mL) is denoted with a dashed line.

**Supplementary Fig. 9**

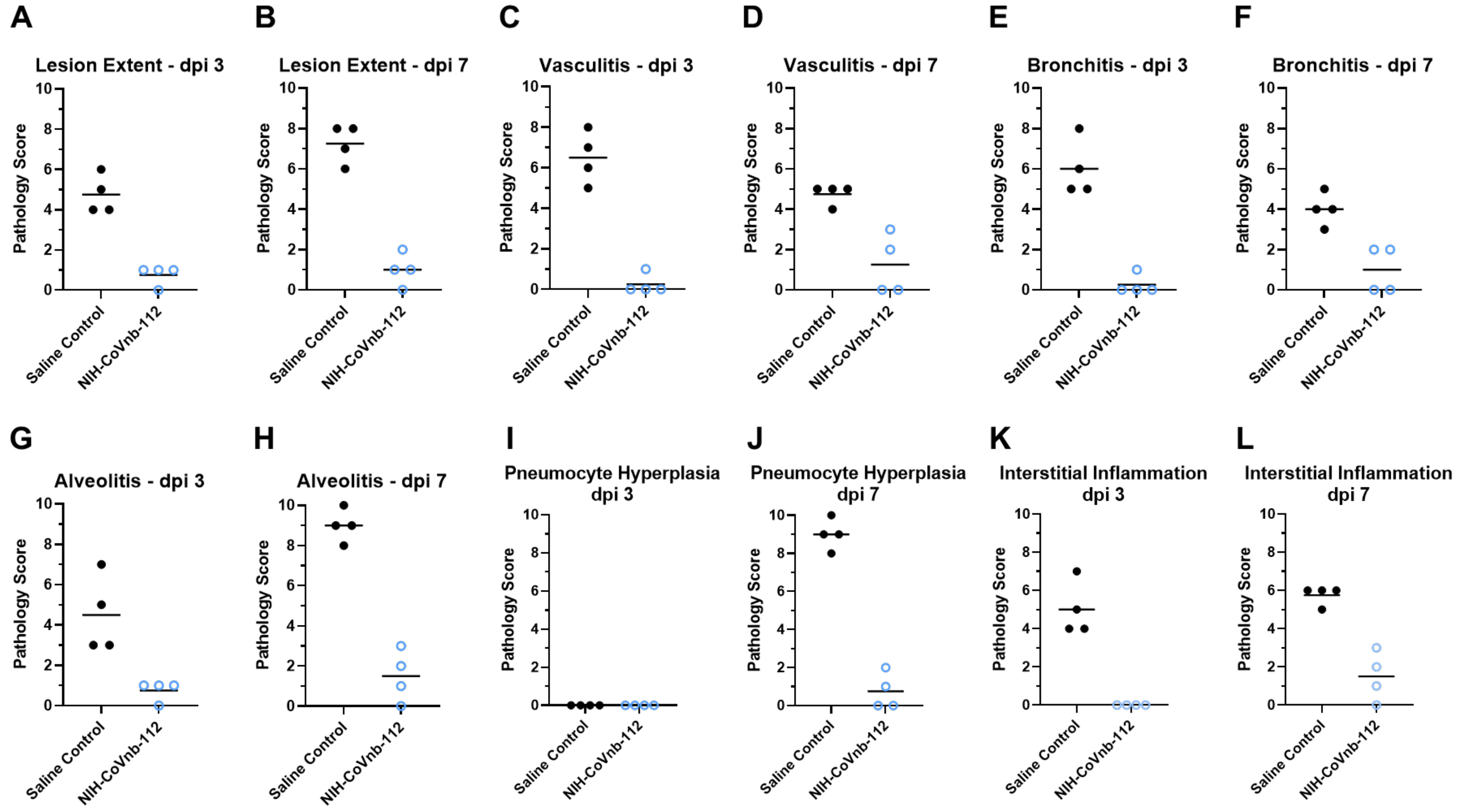

**Supplementary Fig. 9. Histological lung score individual metrics for multi dose nebulization treated hamsters.** Hematoxylin and eosin-stained tissue sections from the day 3 and day 7 post infection animals were semi-quantitively scored by a pathologist blinded to condition using an established scoring metric. Metrics included overall (**a,b**) lesion extent, (**c,d**) vasculitis, (**e,f**) bronchitis, (**g,h**) alveolitis, (**i,j**) pneumocyte hyperplasia, and (**k,l**) interstitial inflammation; each on a 0–4 or 0–5 scale.

**Supplementary Fig. 10**

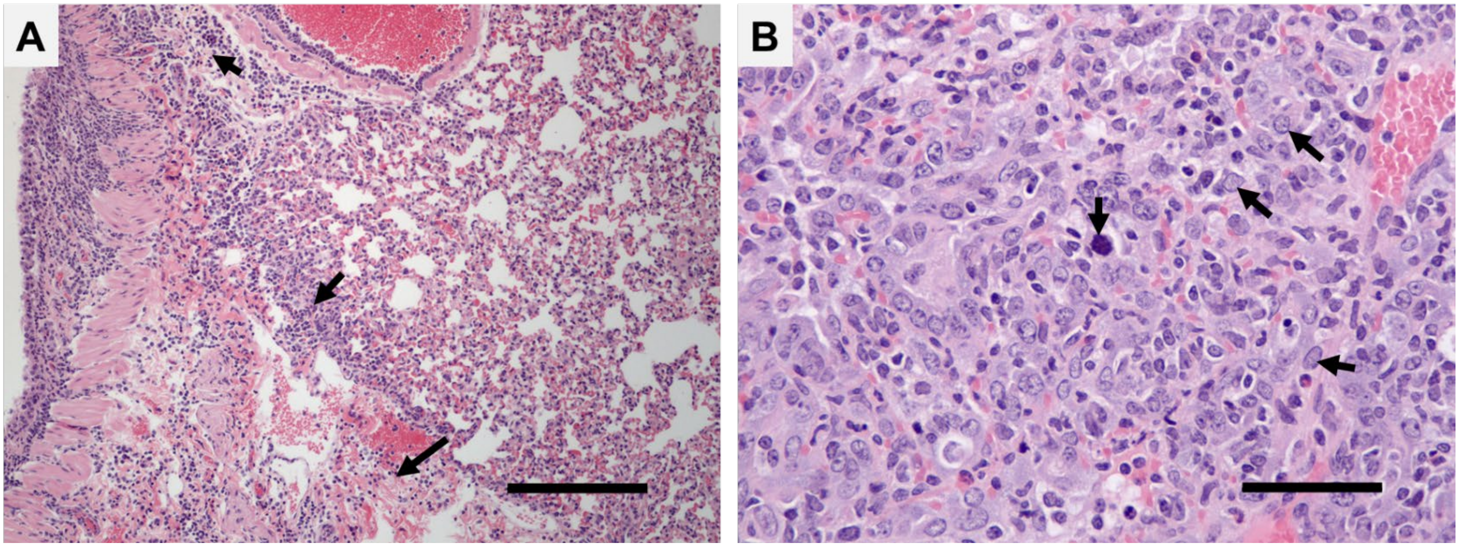

**Supplementary Fig. 10. Interstitial inflammation and pneumocyte hyperplasia pathology in Syrian hamsters following SARS-CoV-2 infection.** (**a**) Representative hematoxylin and eosin-stained lung tissue displaying interstitial inflammation, or thickening of the pulmonary interstitium (arrows), in a saline treated SARS-CoV-2 infected Syrian hamster. Image from a 3dpi animal. Image acquired at 100x magnification. Scale bar = 320μm. (**b**) Representative hematoxylin and eosin-stained lung tissue displaying pneumocyte hyperplasia (single down-facing arrow, note mitosis) and pneumocyte hypertrophy (arrows) in a saline treated SARS-CoV-2 infected Syrian hamster. Image from a 7dpi animal. Image acquired at 400x magnification. Scale bar = 80μm.

### SUPPLEMENTARY REFERENCES

1 Esparza, T. J., Martin, N. P., Anderson, G. P., Goldman, E. R. & Brody, D. L. High affinity nanobodies block SARS-CoV-2 spike receptor binding domain interaction with human angiotensin converting enzyme. *Sci Rep* **10**, 22370, doi:10.1038/s41598-020-79036-0 (2020).

2 Jia, Q. *et al.* Replicating bacterium-vectored vaccine expressing SARS-CoV-2 Membrane and Nucleocapsid proteins protects against severe COVID-19-like disease in hamsters. *NPJ Vaccines* **6**, 47, doi:10.1038/s41541-021-00321-8 (2021).

3 Weiss, M. S. Global indicators of X-ray data quality. *Journal of Applied Crystallography* **34**, 130-135, doi:https://doi.org/10.1107/S0021889800018227 (2001).

4 Karplus, P. A. & Diederichs, K. Linking crystallographic model and data quality. *Science* **336**, 1030-1033, doi:10.1126/science.1218231 (2012).

5 Popov, A. N. & Bourenkov, G. P. Choice of data-collection parameters based on statistic modelling. *Acta Crystallogr D Biol Crystallogr* **59**, 1145-1153, doi:10.1107/s0907444903008163 (2003).

6 Brunger, A. T. Free R value: cross-validation in crystallography. *Methods Enzymol* **277**, 366-396, doi:10.1016/s0076-6879(97)77021-6 (1997).

7 Chen, V. B. *et al.* MolProbity: all-atom structure validation for macromolecular crystallography. *Acta Crystallogr D Biol Crystallogr* **66**, 12-21, doi:10.1107/S0907444909042073 (2010).
